# Development of the Wheat Practical Haplotype Graph Database as a Resource for Genotyping Data Storage and Genotype Imputation

**DOI:** 10.1101/2021.06.10.447944

**Authors:** Katherine W. Jordan, Peter J. Bradbury, Zachary R. Miller, Moses Nyine, Fei He, Max Fraser, Jim Anderson, Esten Mason, Andrew Katz, Stephen Pearce, Arron H. Carter, Samuel Prather, Michael Pumphrey, Jianli Chen, Jason Cook, Shuyu Liu, Jackie C. Rudd, Zhen Wang, Chenggen Chu, Amir M. H. Ibrahim, Jonathan Turkus, Eric Olson, Ragupathi Nagarajan, Brett Carver, Liuling Yan, Ellie Taagen, Mark Sorrells, Brian Ward, Jie Ren, Alina Akhunova, Guihua Bai, Robert Bowden, Jason Fiedler, Justin Faris, Jorge Dubcovsky, Mary Guttieri, Gina Brown-Guedira, Ed Buckler, Jean-Luc Jannink, Eduard D. Akhunov

**Affiliations:** Department of Plant Pathology, Kansas State University, Manhattan, KS, USA; USDA-ARS, Hard Winter Wheat Genetics Research Unit, Manhattan, KS, USA; USDA-ARS, Plant Soil and Nutrition Research Unit, Ithaca, NY, USA; Institute for Genomic Diversity, Cornell University, Ithaca, NY, USA; Department of Agronomy and Plant Genetics, University of Minnesota, St. Paul, MN, USA; Department of Soil and Crop Sciences, Colorado State University, Fort Collins, CO, USA; Department of Crop and Soil Sciences, Washington State University, Pullman, WA, USA; Department of Plant Sciences, University of Idaho, Aberdeen, ID, USA; Department of Plant Sciences and Plant Pathology, Montana State University, Bozeman, MT, USA; Department of Soil and Crop Sciences, Texas A&M AgriLife Reseach, Amarillo, TX, USA; Department of Plan, Soil and Microbial Sciences, Michigan State University, East Lancing, MI, USA; Department of Plant and Soil Sciences, Oklahoma State University, Stillwater, OK, USA; USDA-ARS, Plant Science Research Unit, Raleigh, NC, USA; Integrative Genomics Facility, Kansas State University, Manhattan, KS, USA; USDA-ARS, Cereal Crops Research Unit, Fargo, ND, USA; Department of Plant Sciences, University of California-Davis, Davis, CA, USA

**Author notes:** Corresponding Author: Eduard Akhunov, Department of Plant Pathology, Kansas State University, 1712 Claflin Rd, 4024 Throckmorton Plant Science Center, Manhattan, KS 66506. KWJ is currently affiliated with USDA-ARS, Hard Winter Wheat Genetics Research Unit, Manhattan, KS, USA.

**Keywords:** Wheat, Genotype Imputation, Practical Haplotype Graph, skim-seq, exome capture

## Abstract

To improve the efficiency of high-density genotype data storage and imputation in bread wheat (*Triticum aestivum* L.), we applied the Practical Haplotype Graph (PHG) tool. The wheat PHG database was built using whole-exome capture sequencing data from a diverse set of 65 wheat accessions. Population haplotypes were inferred for the reference genome intervals defined by the boundaries of the high-quality gene models. Missing genotypes in the inference panels, composed of wheat cultivars or recombinant inbred lines genotyped by exome capture, genotyping-by-sequencing (GBS), or whole-genome skim-seq sequencing approaches, were imputed using the wheat PHG database. Though imputation accuracy varied depending on the method of sequencing and coverage depth, we found 93% imputation accuracy with 0.01x sequence coverage, which was only slightly lower than the accuracy obtained using the 0.5x sequence coverage (96.9%). Compared to Beagle, on average, PHG imputation was ~4% (*p-value* = 0.00027) more accurate, and showed 27% higher accuracy at imputing a rare haplotype introgressed from a wild relative into wheat. The reduced accuracy of imputation with GBS data (90.4%) is likely associated with the small overlap between GBS markers and the exome capture dataset, which was used for constructing PHG. The highest imputation accuracy was obtained with exome capture for the wheat D genome, which also showed the highest levels of linkage disequlibrium and proportion of identity-by-descent regions among accessions in our reference panel. We demonstrate that genetic mapping based on genotypes imputed using PHG identifies SNPs with a broader range of effect sizes that together explain a higher proportion of genetic variance for heading date and meiotic crossover rate compared to previous studies.

## Introduction

For the last 10,000 years, intensive selection of bread wheat, *Triticum aestivum*, created varieties adapted to diverse environments and cultivation practices (Balfourier *et al.* 2019; He *et al.* 2019; Walkowiak *et al.* 2020). Recent advances in crop genomics and the availability of reference genomes have accelerated the adoption of sequence-based genotyping technologies for studying the genetics of agronomic traits (Nyine *et al.* 2019) and local adaptation (He *et al.* 2019; Juliana *et al.* 2019, 2020) and facilitated the introduction of genomics-assisted breeding strategies into wheat improvement pipelines (Poland and Rife 2012; Isidro *et al.* 2014). However, the limited genome coverage provided by these genotyping technologies does not support exploration of the entire range of genetic effects conferred by all variants, limiting the utility of the developed genomic diversity and functional genomics resources for understanding genome-to-phenome connections.

The large size (17 Gb) and complexity of the wheat genome present a substantial challenge for sequence-based analysis of genetic diversity. Alignment of short sequence reads to the wheat genome is complicated by high levels of sequence redundancy resulting from two rounds of recent whole genome duplication (IWGSC, 2018), and the recent propagation of transposable elements (TEs) comprising nearly 90% of the genome (Wicker *et al.* 2018). Therefore, the efforts of the wheat research community were focused primarily on sequencing complexity-reduced genomic libraries produced by either enzymatic digests or by targeted sequence capture. These efforts have resulted in a detailed description of the population-scale haplotypic diversity in the low-copy genomic regions in large sets of genetically and geographically diverse wheat lines and breeding populations (He *et al.* 2019; Juliana *et al.* 2019; Pont *et al.* 2019). While these resources have been useful for genotype imputation in populations genotyped using either SNP-based arrays or genotyping-by-sequencing (GBS) methods (Jordan *et al.* 2015; Shi *et al.* 2017; Juliana *et al.* 2019; Nyine *et al.* 2019), the relatively small number of shared markers between the reference and inference populations limits the number of imputed genotypes, thus diminishing the utility of genotype imputation in wheat genetic studies and breeding.

High-quality reference genomes and a reduction in the cost of sequencing presented opportunities for the characterization of genetic diversity by direct sequencing of either whole genomes or genomic regions targeted by sequence capture (Malmberg *et al.* 2018; He *et al.* 2019; Walkowiak *et al.* 2020). While these sequence-based genotyping approaches generate unbiased information about the genetic variants of various frequency classes and genomic locations, large-scale population sequencing of species with large genomes, including many important agricultural crops, remains costly. This issue has been addressed by combining low-coverage sequencing of whole genomes with the prediction of missing genotypes using imputation tools, thereby increasing the power of association mapping and facilitating the detection of causal variants (Davies *et al.* 2016; Das *et al.* 2018; Rubinacci *et al.* 2021).

Recently, a novel strategy referred to as Practical Haplotype Graph (PHG), was proposed to improve the efficiency of sequence-based genotyping data storage and imputing genotypes in low-coverage sequencing datasets (Jensen *et al.* 2020; Valdes Franco *et al.* 2020). The PHG is capable of storing genotyping data generated using diverse genotyping technologies as a graph of haplotypes of founder lines and is used for predicting missing genotypes in populations characterized by various sequence- or array-based genotyping strategies. By reducing the constraints associated with large-scale genotyping data storage, processing, and utilization, this tool is another step towards leveraging the existing community-generated genomic diversity resources in breeding and research applications. We used skim-seq, whole-exome capture, genotyping-by-sequencing, and array-based genotyping datasets generated by the USDA-NIFA WheatCAP to develop a wheat PHG database and evaluate its performance for genotype imputation in wheat lines of different levels of relatedness and different depths of genome coverage.

## Methods

### Library prep

DNA was extracted from two-week old leaf tissue of germinated seedlings grown under greenhouse conditions from breeding programs across the United States (Table S1). DNA was extracted using Qiagen DNeasy kit following the manufacturer’s protocol. DNA was quantified with Picogreen (Sage Scientific) and wheat exome capture was performed on each sample targeting the non-redundant low-copy portion of the genome. Briefly, wheat exome captures designed in collaboration with Nimblegen targeted 170 Mb of sequence covering about 80,000 transcripts (Krasileva *et al.* 2017). The barcoded genomic libraries were pooled at 12- or 96-plex levels, and sequenced on NextSeq (Kansas State University Integrated Genomics Facility) and NovaSeq (Kansas University Medical Center) platforms using 2 × 150 bp read runs to produce sequence data providing about 30x coverage of the exome capture target space.

Genomic libraries for low-coverage sequencing were prepared for 18 samples from the NAM18 family (Jordan *et al.* 2018) using Illumina DNA Prep Kit along with the Illumina’s Nextera CD adapters. Sequencing was performed on the Illumina NextSeq platform to produce ~0.1x coverage.

### Data processing

The quality of sequence reads was assessed using NGSQC toolkit v.2.3.3 (Patel and Jain 2012). The sequence reads were aligned to the wheat reference genome RefSeq v.1.1 (IWGSC, 2018) using HiSat2 (Kim *et al.* 2015) retaining only uniquely mapped reads. The resulting alignments were processed using the GATK pipeline (McKenna *et al.* 2010) to generate a genome variant call file (g.vcf) for each accession.

The raw variant calls generated by GATK for exome capture data were filtered using *bcftools* (Danecek *et al.* 2021) to retain variants with minor allele frequency ≥ 0.015 and missing data < 10%. Filtered GATK variants were combined with 90K genotypic data (Wang *et al.* 2014), producing high quality filtered variants (henceforth, HQ-SNPs) that were used for assessing the accuracy of the PHG-based imputation.

### Wheat PHG database construction

The Wheat PHG database was built using PHG version 0.017. Instructions for creating the PHG along with source code are located with the PHG wiki: https://bitbucket.org/bucklerlab/practicalhaplotypegraph/wiki/Home. The approaches and parameters for constructing the Wheat PHG were discussed and developed during two PHG workshops organized at Cornell University. The first step of the PHG database construction is to create reference ranges for data storage and variant imputation (Fig. S1). In this case, “informative” reference ranges were chosen by extending the high confidence gene model coordinates from Chinese Spring RefSeq v.1.1 (IWGSC, 2018) 500 bp in each direction. Adjacent ranges were merged if the boundaries lie within 500 bp from each other. This resulted in a final set of 106,484 informative reference ranges across the genome from the Chinese Spring accession, while the rest of the genome was deemed non-informative and represents intergenic ranges across the genome of Chinese Spring (Fig. S1).

The second step in the PHG pipeline populates the database with sequence data from diverse accessions across the reference ranges (Fig. S1). Pre-processed exome capture g.vcf files for 65 accessions, including 58 *Tricitum aestivum* accessions, 3 *Aegilops tauschii* accessions, 3 *Triticum turgidum* subsp. *durum* wheat cultivars, and one *Triticum turgidum* subsp. *dicoccum* accession (Table S1) generated by GATK (McKenna *et al.* 2010) were loaded into the PHG, creating a database of 6,705,472 haplotypes, which is representative of the diversity across the wheat breeding programs within the US and breeding lines from the Great Plains region.

The third step of the PHG pipeline is to create consensus haplotypes for the reference ranges, using the sequence information of the 65 accessions (Fig. S1). This step collapses the raw haplotypes into consensus haplotypes using a user-defined maximum divergence (mxDiv) parameter set to 0.0001. This translates into the clustering of raw haplotypes that contain less than 1 bp divergence per 10,000 bp into a common haplotype. The value of the mxDiv parameter was based on prior diversity estimates in wheat (Akhunov *et al.* 2010; Jordan *et al.* 2015), and aimed at retaining a manageable number of haplotypes per reference range as described in Jensen *et al*. (2020). In addition to the mxDiv parameter, we set minTaxa = 1, which retains haplotypes present in only one accession and facilitates the imputation of rare haplotypes. Using these parameters, a total of 712,733 consensus haplotypes were detected, which is approximately 6.7 haplotypes per informative reference range, similar to ~5 haplotypes per reference range reported in the sorghum PHG (Jensen *et al.* 2020).

At the imputation step, the low coverage sequence data were aligned to the consensus haplotypes stored in the PHG database (Fig S1), and a Hidden Markov model was used to infer the paths through the practical haplotype graph that match the mapped reads while determining the missing haplotypes. The variants were imputed using the haplotype structure stored in the database, and exported as a vcf file. By using minReads = 0 parameter, variant calls were imputed for all variable positions in the wheat PHG database.

To assess the effect of genome coverage depth on imputation accuracy, we used *seqtk* (Li 2012) to generate down-sampled datasets from the 170Mb wheat exome capture data representative of 0.01x (5,667 paired-end (PE) reads per accession), 0.1x (56,667 PE per accession), and 0.5x (283,333 PE reads per accession) depth of coverage for 20 breeding lines from the US Great Plains (Table S1). This set of 20 breeding lines included four lines (Duster, Overley, NuPlains, and Zenda), which were used to build the PHG database.

In addition, we assessed the accuracy of imputation in genotyping datasets generated using GBS of genomic libraries prepared from MseI-PstI digested DNA (Saintenac *et al.* 2013) and whole-genome skim sequencing (Malmberg *et al.* 2018). We used previously published GBS data produced for 75 recombinant inbred lines from the wheat nested-association mapping (NAM) population (Jordan *et al.* 2018) that included an average of 1.85 million (1x 100bp) reads per accession. In the current study, we performed the whole-genome skim sequencing on a set of 18 recombinant inbred lines, using 2 × 150 bp sequencing reads providing on average 6.1 million paired-end reads per accession, which represents ~0.1x genome coverage (Table S2).

### PHG Imputation Accuracy

The accuracy of genotype calls for each accession was determined by dividing the number of matching genotype calls between the HQ-SNPs and the PHG-imputed SNP data by the total number of overlapped genotype calls. For down-sampled datasets generated from the exome capture data, imputation accuracy was estimated using nearly 400,000 genotype calls per accession at each sequence coverage level. Imputation accuracy comparisons by genome, and by MAF category were performed using ANOVA from *car* and *lme4* R packages. The imputation accuracy estimates for the GBS and whole-genome skim-sequencing data were based on approximately 5,000 HQ-SNPs genotype calls per accession.

### Comparison of accuracy between Beagle v.5.0 and PHG-based genotype imputation

The accuracy of PHG genotype imputation was compared to the accuracy of imputation with Beagle v.5.0. (Browning and Browning 2013). For this purpose, we used the *reference panel* of 65 accessions that was also utilized to construct the wheat PHG. The genotyping data for a *target panel* were generated by calling genotypes using the down-sampled sequence reads following the same SNP calling procedures described above. The genotype calls were produced for the 20 Great Plains accessions at each genome coverage level. Imputation was performed using Beagle v.5.0 with the default parameters. The genotype calls imputed with Beagle were compared to the HQ-SNP dataset (see above) to assess the overall concordance and concordance of minor allele calls. On average, the estimates of accuracy were based on about 323,000 genotype calls per accession. Formal comparisons of the imputation accuracy between Beagle v5.0 and PHG imputation methods by coverage level for 0.01x and 0.1x were performed using paired t-tests in R. At each coverage level, imputation using PHG was statistically more accurate (0.01x: *p-value* = 2.7 × 10^−4^; 0.1x: *p-value* = 2.2× 10^−7^).

### Diversity analysis

Diversity statistics (π and Tajima’s D) were estimated using TASSEL v5.2.65 (Bradbury *et al.* 2007) in sliding windows of 2,000 SNPs per window stepping 1,000 SNPs at a time, and mean values per genome were calculated. The identity-by-descent (IBD) analysis was determined using Beagle v.4.1 with the default parameters (Browning and Browning 2013), and considered to be significant at LOD ≥ 3.0. Overlap between the IBD segments was determined using the MultiIntersectBed tool of the Bedtools suite v.2.26.0 (Quinlan and Hall 2010). Pairwise linkage disequilibrium (LD) was determined using PLINK v.1.90b3.45 (Purcell *et al.* 2007) by calculating the coefficients of determination (*r*^2^) for all possible pairwise combinations of SNP sites on the same chromosomes.

### Stepwise regression using the PHG imputed markers

The parental lines of a family of 75 recombinant inbred lines (RILs) from the spring wheat NAM panel (Jordan *et al.* 2018) were included into the panel of 65 accessions that were used to construct the wheat PHG. We ran PHG imputation on the GBS data generated for 75 RILs, and imputed genotypes for 1.457 million sites. These sites were filtered to retain variants that segregate between the parental lines, and have allele frequencies between 0.35-0.65 in the RIL population. These variants were subsequently thinned using PLINK (Purcell *et al.* 2007) to remove markers that had an *r*^2^ ≥ 0.6 within a 50 SNP window, stepping 10 SNPs at a time. The resulting set of 9,806 markers with no missing data was used for stepwise regression mapping performed with the ICIM software v.4.1.0.0 (Meng *et al.* 2015) with markers entering and exiting the model with *p-value* < 0.0001. The estimates of the Total number of CrossOvers (TCO) and the distal CrossOvers (dCO) were taken from the previous analyses of the spring wheat NAM population for family NAM1 (Jordan *et al.* 2018). Heading dates were measured in three locations for two growing seasons (Montana, South Dakota, Washington) for the 75 RILs and three checks. Best linear unbiased predicitions (BLUPs) for each line were estimated using the following linear mixed model with *lmer* package in R:

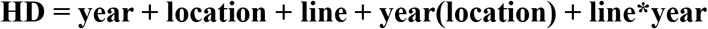

where location, year, and location nested within year are fixed variables, and the line and line-by-year interaction terms are random variables.

## Results

### The Wheat PHG database development

A wheat PHG database was created using whole-exome capture data from a set of 65 wheat accessions (Table S1) contributed by the major U.S. wheat breeding programs, as well as the parental lines used for the genetic analyses of the yield component traits in WheatCAP (www.triticeaecap.org). This set of accessions was selected from a larger diversity panel of nearly 250 wheat cultivars assembled in coordination with the U.S. wheat breeding programs to build a genomic resource to be used as a reference panel for genotype imputation. This diverse set of 65 accessions is comprised of mostly spring and winter bread wheat cultivars, but it also included three accessions of the diploid ancestor of the wheat D genome, *Aegilops tauschii* (accessions TA1615, TA1718, and TA1662/PI603230), and four accessions of tetraploid wheat (three *Triticum turgidum* subsp. *durum* wheat cultivars Langdon, Ben, and Mountrail and one domesticated emmer, *Triticum turgidum* subsp. *dicoccum*, accession PI41025).

For constructing the wheat PHG, the wheat genome was split into a set of informative reference ranges that represent the high confidence gene models in the IWGSC RefSeq v.1.1 (IWGSC, 2018). By using the predicted gene models to define reference ranges, we aimed to reduce the impact of erroneous genotype calling associated with the misalignments of sequence reads to the repetitive portion of the wheat genome (Wicker *et al.* 2018) on the estimation of linkage disequilibrium (LD) and detecting haplotype blocks. A total of 106,484 reference ranges spanning all 21 chromosomes were defined (Fig S1; Table S3), with an average of 5,070 reference ranges per chromosome; chromosome 4D contains the lowest (3,612 ranges) and chromosome 2B harbors the highest (6,221 ranges) number of reference ranges.

Using the 65 accessions to populate the wheat PHG database, we discovered 1,473,670 variants across the 106,484 reference ranges, of which 1,457,321 are high quality, bi-allelic SNPs (Table S3). The inclusion of three diploid *Ae. tauschii* accessions into the panel increased the number of variable sites detected in the D genome lineage, which is the least polymorphic genome in bread wheat (Wang *et al.* 2013; Jordan *et al.* 2015; He *et al.* 2019). Excluding the variants from *Ae. tauschii*, we found that 161,226 (31%) sites in the D genome were monomorphic among the bread wheat cultivars. Similarly, we found that 31,486 SNPs (7%) in the A genome and 32,228 SNPs (6%) in the B genome are contributed by the domesticated emmer and durum lines, and are monomorphic in hexaploid wheat. These private SNPs explain the high levels of divergence between the domesticated emmer and *Ae. tauschii* accessions from the hexaploid wheat lines (Fig. 1a). The overall patterns of genetic diversity and allele frequency distribution in the D genome compared to those in the A and B genomes were consistent with the population bottleneck (Table 1): 1) diversity mean estimates for the D genome were less than 2.3-fold that of the A and B genomes, (π_D_ = 0.076, π_A_ = 0.175, and π_B_ = 0.182; Table 1), 2) the estimates of Tajima’s D were lower in the D genome than in the A and B genomes (Tajima’s D_D_= −2.19, Tajima’s D_A_= −0.67, and Tajima’s D_B_ = −0.55, Table 1), 3) the mean minor allele frequencies (MAF) were greater in the A and B genomes than in the D genome (MAF_A_= 0.12, MAF_B_= 0.12, and MAF_D_= 0.05), and 4) LD drops to half of its initial value (*r*^2^ ≤ 0.33) at 20 Mb in the D genome, whereas in the A and B genomes LD drops to the same level at 12 and 10 Mb, respectively (Table 1, Figure 1b).

**Figure 1.**
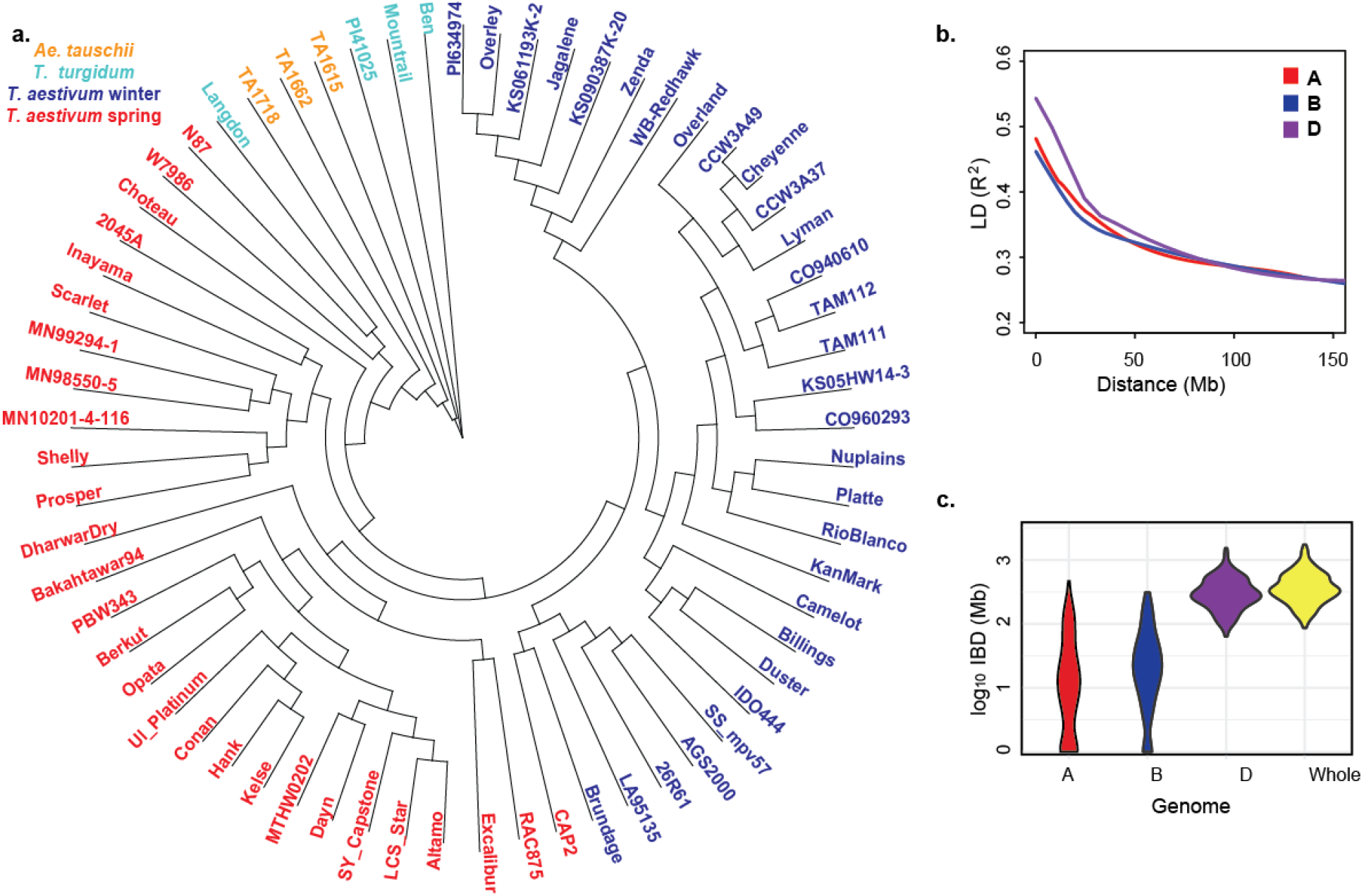
Genetic diversity of 65 accessions of wheat and its diploid and tetraploid relatives used for developing the Wheat PHG. **a.** Neighbor-joining tree of accessions used for constructing the Wheat PHG. **b.** The rate of LD decay in the A, B and D genomes of wheat. **c.** The length of pair-wise IBD between the parental lines from different breeding programs used in WheatCAP.

**Table 1.** Estimates of genetic diversity (π), minor allele frequency (MAF), Tajima’s D and linkage disequilibrium in the population used for constructing the Wheat PHG.

| Diversity statistic | A genome | B genome | D genome |
| --- | --- | --- | --- |
| No. SNPs | 430,050 | 504,260 | 523,011 |
| MAF | 0.116 | 0.122 | 0.050 |
| $\pi$ (per bp) | 0.175 | 0.182 | 0.076 |
| Tajima's D | -0.673 | -0.552 | -2.192 |
| LD* ($r^2 \leq 0.33$) | 12.2 Mb | 9.8 Mb | 20.0 Mb |

\*distance at which LD drops to half of its initial value ( $r^2 \leq 0.33$ ).

The accuracy and the rate of genotype imputation are affected by the proportion of shared genetic ancestry among individuals in a population (Browning and Browning 2013). For each WheatCAP parental line included in the Wheat PHG, we estimated the length of genomic segments sharing identity-by-descent (IBD) with other lines in the panel. On average, the pairs of parents had 451 Mb (~3%) of IBD segments (Table S4), suggesting distant relationships among the WheatCAP parental lines. However, the estimates of the total length of IBD segments among cultivars were quite variable (Figure 1c). For example, in cultivars Prosper from North Dakota and Shelly from Minnesota, the length of shared IBD segments was nearly 1.29 Gb (8.6%), whereas hard winter wheat cultivars Lyman (South Dakota) and Overley (Kansas) shared only 128 Mb (0.85%) of IBD segments. The average length of IBD segments shared by the distantly related durum wheat and domesticated emmer parents was only 57.6 Mb. Across all breeding programs, we detected 556 regions sharing IBD, with an average IBD segment length of 12.2 Mb. Over half (53%) of the IBD segments overlapped with a segment from at least one other breeding program, translating to more than 1.68 Gb of the genome shared between any two wheat breeding programs. This estimate includes 1.49 Gb of shared IBD in the D genome (89%), while only 86.4 Mb and 105.7 Mb of IBD with other breeding programs were detected in the A and B genomes, respectively. The genomic segments sharing IBD with most of the wheat lines were located on chromosomes 7D (568 Mb - 571 Mb) and 3D (496.6 Mb - 505 Mb), which were common to seven breeding programs.

In addition to the WheatCAP lines, we selected 21 hard red winter wheat cultivars from the U.S. Great Plains for constructing the PHG database (Table S1). Pairwise comparisons among these lines showed that, on average, they share 416 Mb of IBD segments, with an average IBD segment length of 13 Mb, and nearly 83% of all shared IBD regions are located in the D genome (Table S5). This finding is consistent with the lack of diversity among breeding lines in the D genome (Chao *et al.* 2010) and the high levels of shared ancestry among the lines from the U.S. Great Plains’ breeding programs.

### Genotype imputation using the Wheat PHG

We used several low-coverage sequencing datasets to assess the imputation performance of the wheat PHG. First, we used 20 spring and winter wheat lines (Table S1) from the U.S. wheat breeding programs sequenced using the whole-exome capture approach (Krasileva *et al.* 2017; He *et al.* 2019) to mimic a low-coverage sequencing experiment. We down-sampled the raw unmapped Illumina paired-end reads generated for each accession to create datasets with three levels of sequence coverage depths (0.01x, 0.1x, and 0.5x) for the regions targeted by the exome capture assay. The accuracy of imputation achieved using the Wheat PHG was estimated by comparing the concordance of imputed genotype calls with the genotype calls from the HQ-SNP set generated using the 90K iSelect array (Wang *et al.* 2014) and the high-coverage (20-30x coverage) exome sequencing.

On average, using 0.5x coverage down-sampled exome capture data, we achieved 96.9% imputation accuracy, ranging from 95% to 98% among lines (Figure 2a, Table 3). Five- and fifty-fold reduction in the depth of read coverage for the inference panel did not result in a substantial reduction in the accuracy of imputation. The mean accuracy of PHG imputation was 96% (94-98% range) with 0.1x coverage depth, and 93% (91-98% range) with as little as 0.01x coverage depth (Figure 2a, Tables 2 and 3). These results suggest that the imputation method in the PHG could effectively use 0.01x exome coverage data to adequately capture the haplotypic diversity of the inference panel to achieve 93% imputation accuracy. The imputation accuracy varied among the wheat genomes, likely due to genome-specific differences in the extent of LD and haplotypic diversity (Jordan *et al.* 2015). At 0.01x coverage depth, the accuracy of genotype imputation in the D genome was 95.5%, which was 3.2% and 4.3% more accurate (*p-value* _(ANOVA)_= 3.73×10^−5^) than imputation in the A (92.3%), and the B genomes (91.2%), respectively (Table 2; Figure 2b). The higher extent of LD in the D genome appears to contribute to more accurate genotype imputation compared to that in the A and B genomes, which show faster rates of LD decay and lower proportions of the genome sharing IBD segments in the panel used to build the PHG database.

**Figure 2.**
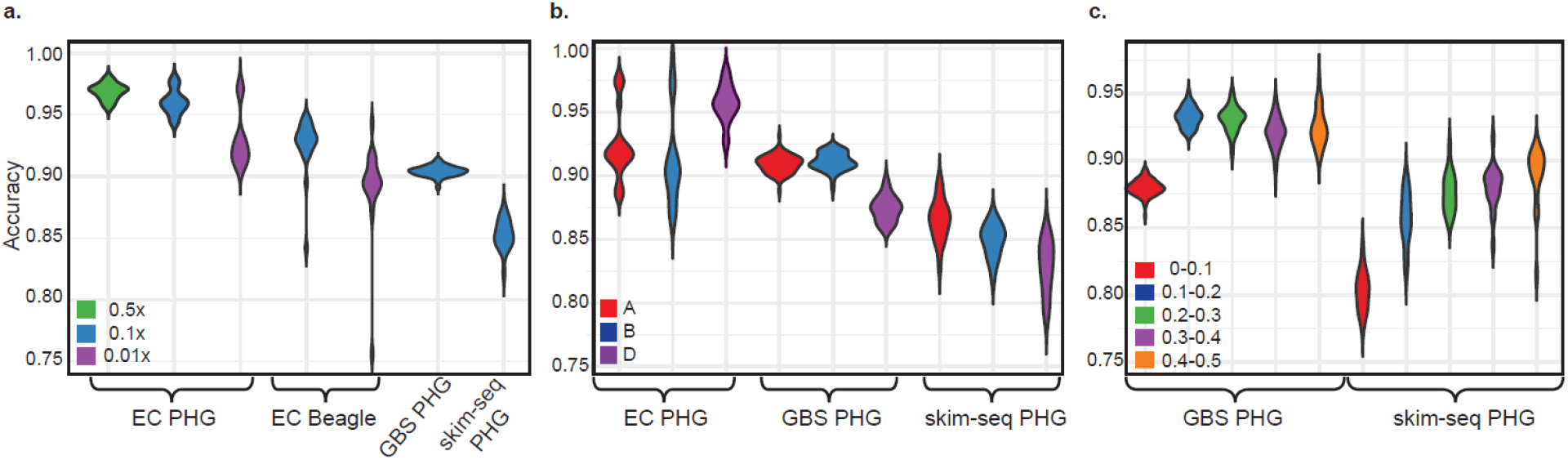
The accuracy of imputation using the wheat PHG. **a.** The impact of sequence coverage and the method of imputation on accuracy, (EC: n=20; GBS: n=75; skim-seq: n=18) **b.** Accuracy of imputation in the A, B and D genomes of wheat using exome capture (EC), GBS and whole genome skim-seq data. **c.** Accuracy of imputation for alleles with different minor allele frequency.

**Table 2.** The accuracy of PHG imputation in different wheat genomes.

| Wheat genomes | Exome capture* (0.01x), % | GBS*, % | skim-seq* (0.1x), % |
| --- | --- | --- | --- |
| **Total** | 92.6 | 90.4 | 85.3 |
| **A** | 92.3 | 90.9 | 86.5 |
| **B** | 91.2 | 91.1 | 84.9 |
| **D** | 95.5 | 87.4 | 82.9 |

\*Accuracies by approach are comprised of different germplasm, EC: n=20, GBS: n=75, skim-seq: n=18

**Table 3.**
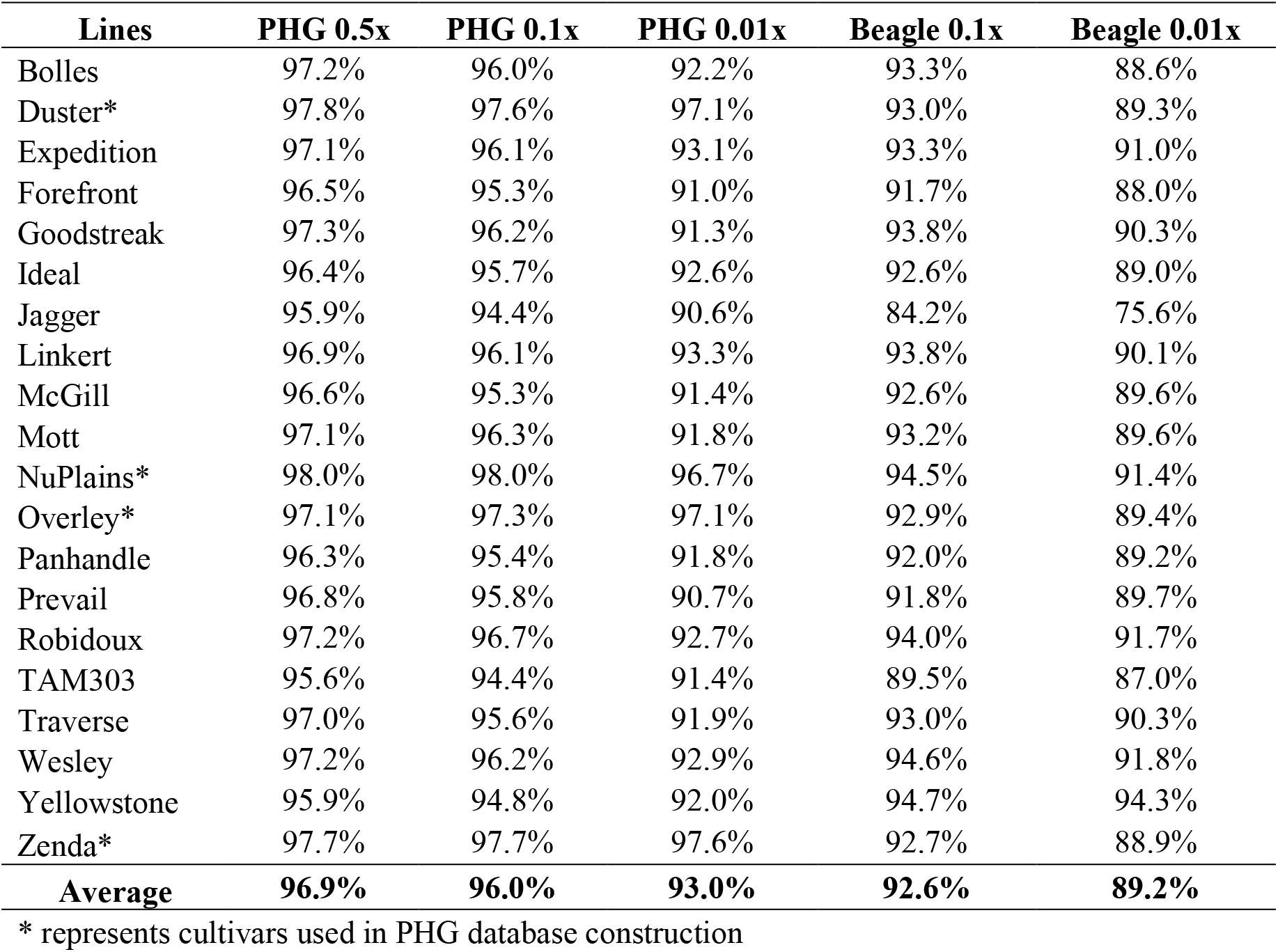
Comparison of imputation accuracy between PHG and Beagle using exome capture data.

| Lines | PHG 0.5x | PHG 0.1x | PHG 0.01x | Beagle 0.1x | Beagle 0.01x |
| --- | --- | --- | --- | --- | --- |
| Bolles | 97.2% | 96.0% | 92.2% | 93.3% | 88.6% |
| Duster* | 97.8% | 97.6% | 97.1% | 93.0% | 89.3% |
| Expedition | 97.1% | 96.1% | 93.1% | 93.3% | 91.0% |
| Forefront | 96.5% | 95.3% | 91.0% | 91.7% | 88.0% |
| Goodstreak | 97.3% | 96.2% | 91.3% | 93.8% | 90.3% |
| Ideal | 96.4% | 95.7% | 92.6% | 92.6% | 89.0% |
| Jagger | 95.9% | 94.4% | 90.6% | 84.2% | 75.6% |
| Linkert | 96.9% | 96.1% | 93.3% | 93.8% | 90.1% |
| McGill | 96.6% | 95.3% | 91.4% | 92.6% | 89.6% |
| Mott | 97.1% | 96.3% | 91.8% | 93.2% | 89.6% |
| NuPlains* | 98.0% | 98.0% | 96.7% | 94.5% | 91.4% |
| Overley* | 97.1% | 97.3% | 97.1% | 92.9% | 89.4% |
| Panhandle | 96.3% | 95.4% | 91.8% | 92.0% | 89.2% |
| Prevail | 96.8% | 95.8% | 90.7% | 91.8% | 89.7% |
| Robidoux | 97.2% | 96.7% | 92.7% | 94.0% | 91.7% |
| TAM303 | 95.6% | 94.4% | 91.4% | 89.5% | 87.0% |
| Traverse | 97.0% | 95.6% | 91.9% | 93.0% | 90.3% |
| Wesley | 97.2% | 96.2% | 92.9% | 94.6% | 91.8% |
| Yellowstone | 95.9% | 94.8% | 92.0% | 94.7% | 94.3% |
| Zenda* | 97.7% | 97.7% | 97.6% | 92.7% | 88.9% |
| **Average** | **96.9%** | **96.0%** | **93.0%** | **92.6%** | **89.2%** |

\* represents cultivars used in PHG database construction

We compared the performance of the wheat PHG to one of the commonly used low-coverage imputation methods implemented in Beagle v5.0 (Browning and Browning 2013). For this purpose, the panel of 65 accessions included into the wheat PHG database was used as the reference panel, and an independent set of 20 wheat cultivars from the U.S. wheat breeding programs was used as the inference panel. Overall, Beagle imputed missing genotypes with 89.2% accuracy for this set of 20 lines at 0.01x coverage (ranging from 76% to 94%), and 92.6% (ranging from 84% to 95%) at 0.1x coverage (Figure 2a, Table 3). Direct comparisons of imputation methods show PHG imputation statistically outperformed Beagle imputation by 4% at both coverage levels (*p-value* _0.1x (t-test)_ = 2.0×10^−7^; *p-value* _0.01x (t-test)_ = 2.7×10^−4^).

We conducted the analyses of PHG performance in the datasets down-sampled from exome capture data generated for four cultivars Duster, Overley, NuPlains, and Zenda, that were included in the wheat PHG construction. The accuracy of PHG-based imputation for these four cultivars was statistically higher (ANOVA for different levels of sequence coverage: *p-value* _0.5x_ = 0.004; *p-value* _0.1x_ = 2.0 × 10^−5^; *p-value* _0.1x_ = 7.4 × 10^−10^) than for other cultivars at all levels of sequence coverage (Fig. S2a). No similar relationship between the presence of specific haplotypes in the reference panel and imputation accuracy was observed for Beagle. We further explored this relationship by analyzing genotype imputation results in cultivar Jagger, which showed a substantial reduction in imputation accuracy in the low sequence coverage datasets (0.1x and 0.01x coverage) imputed using Beagle (Fig. S2a). We assumed that one of the likely factors contributing to the decreased imputation performance of Beagle in cultivar Jagger was the presence of wild-relative introgression from *Ae. ventricosa* on chromosome 2A (Cruz et al. 2016). Because cultivar Overley, which was used to build the PHG database, also carries this *Ae. ventricosa* introgression (Cruz et al. 2016), we could evaluate the impact of the presence of the rare introgressed haplotype in both the PHG database and the Beagle’s reference panel on imputation accuracy. The chromosome-by-chromosome assessment of imputation accuracy for cv. Jagger in the 0.01x coverage dataset showed modest accuracy (90%) for chromosome 2A using PHG. However, for the same chromosome, the imputation accuracy of Beagle reached only 63% (Fig. S2b). The accuracy of Beagle imputation was also low for other chromosomes (2D, 6A, 7A) (Fig. S2b), which suggests that cv. Jagger likely carries other regions with unique haplotypes (Kippes *et al.* 2018; Walkowiak *et al.* 2020) poorly represented in the reference set used for Beagle imputation. For the same three chromosomes, the accuracy of PHG imputation was higher than that obtained using Beagle, indicating that PHG is more effective at utilizing the rare haplotypes in the reference panel for imputation than Beagle.

### Imputation accuracy with reduced coverage sequencing data

To this point, we tested the imputation accuracy using the same type of genomic data (whole-exome capture) as was used to populate the database. We also evaluated the utility of the developed PHG database for imputing genotypes in the inference panels genotyped using two cost-effective complexity-reduced sequencing approaches, genotyping-by-sequencing (GBS) (Elshire *et al.* 2011; Saintenac *et al.* 2013) and whole-genome skim-seq (Malmberg *et al.* 2018). First, utilizing GBS reads generated for a set of recombinant inbred lines (RILs) from the spring wheat NAM panel (Jordan *et al.* 2018), we performed genotype imputation at 1.4 million variable sites. The parents of these NAM RILs were included into the wheat PHG construction. The mean accuracy of imputation across the 75 RILs was 90.4%, ranging from 89 - 91.4% across individual lines (Figure 2a, Table S6). These estimates of accuracy were only slightly lower than those observed for the imputed genotypes in the down-sampled exome capture data, and likely explained by the relatively small overlap (~5%) between the sites in the GBS and exome capture datasets (Jordan *et al.* 2015). Overall, this result indicates that the PHG based on the panel of wheat lines re-sequenced by exome capture assay provides accurate imputation on the inference population characterized by complexity-reduced sequencing approaches similar to the GBS method.

We also evaluated the wheat PHG in a set of NAM RILs genotyped using the whole-genome skim-seq approach. The genomic libraries generated for a set of RILs from the spring wheat NAM population (Jordan *et al.* 2018; Blake *et al.* 2019) were sequenced on an Illumina sequencer (2 × 150 bp run) to provide ~0.1x genome coverage. The accuracy of PHG-imputed genotypes in the skim-seq dataset (85.3%) was lower than that obtained for genotypes in either the exome capture or GBS datasets. This lower accuracy likely is associated with a lower proportion of skim-seq reads, mostly represented by reads from the repetitive regions, uniquely mapped to the wheat genome compared to the proportion of uniquely mapped reads from the exome capture and GBS datasets, which are enriched for the low-copy genomic regions (Saintenac *et al.* 2013; Jordan *et al.* 2015). The accuracy of imputation varied across different SNP frequency classes. For SNPs with MAF > 0.1, the accuracy of imputation improved by at least 5% for both GBS and skim-seq genotypes. The accuracy reached nearly 90% for skim-seq and 93% for GBS datasets when the MAF were ≥ 0.2 (Table 4, Figure 2c).

**Table 4.**
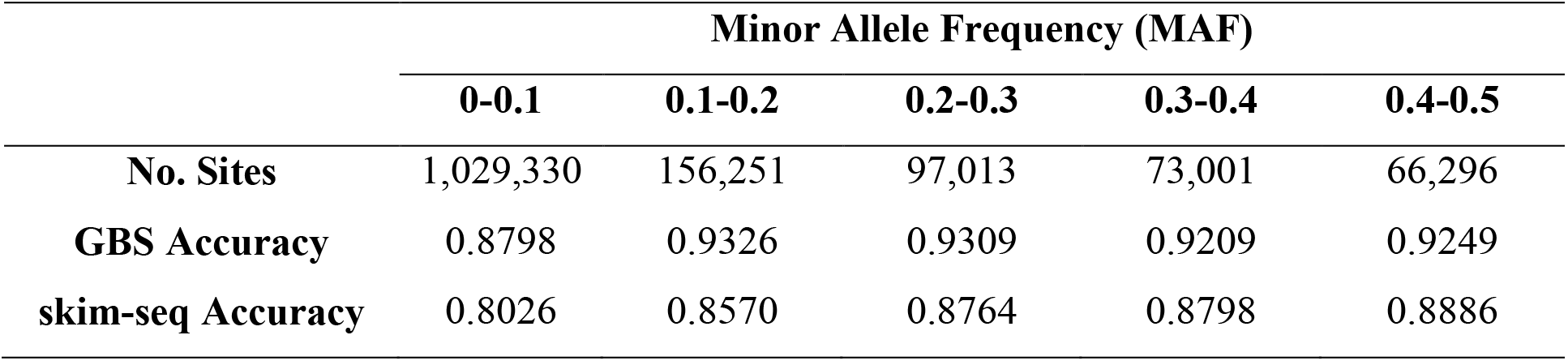
Relationship between minor allele frequency and the accuracy of imputation.

### Genetic analyses of trait variation using the imputed genotypes

The ability to accurately impute genotypes across the genome in low-coverage sequencing datasets provides a cost-effective means for advancing the genetic dissection of trait variation. We used the imputed genotypes to assess the genetic contribution to heading date (HD) variation in the nested association mapping (NAM) family previously used for studying the genetics of recombination rate variation in wheat (Jordan *et al.* 2018). A stepwise regression (SR) was applied to identify variants associated with phenotypic variation. Before mapping, co-segregating redundant markers were removed, resulting in nearly 10,000 markers with no missing data. The SR method identified 11 SNPs together explaining 90% of the variance in heading date, which was measured over two years at three locations (Fig 3, Table S7). Among these SNPs are loci with modest effect sizes located on the long arms of chromosomes 5A and 5D, within 10 Mb from the *Vrn-A1* and *Vrn-D1* loci, which play a major role in the regulation of flowering in wheat (Distelfeld *et al.* 2009). In addition, significant SNPs on chromosomes 1B and 1D were mapped to the regions within 50 Mb of the *Elf-3* gene, which is associated with the transition from vegetative to reproductive growth in wheat (Alvarez *et al.* 2016; Zikhali *et al.* 2016).

**Figure 3.**
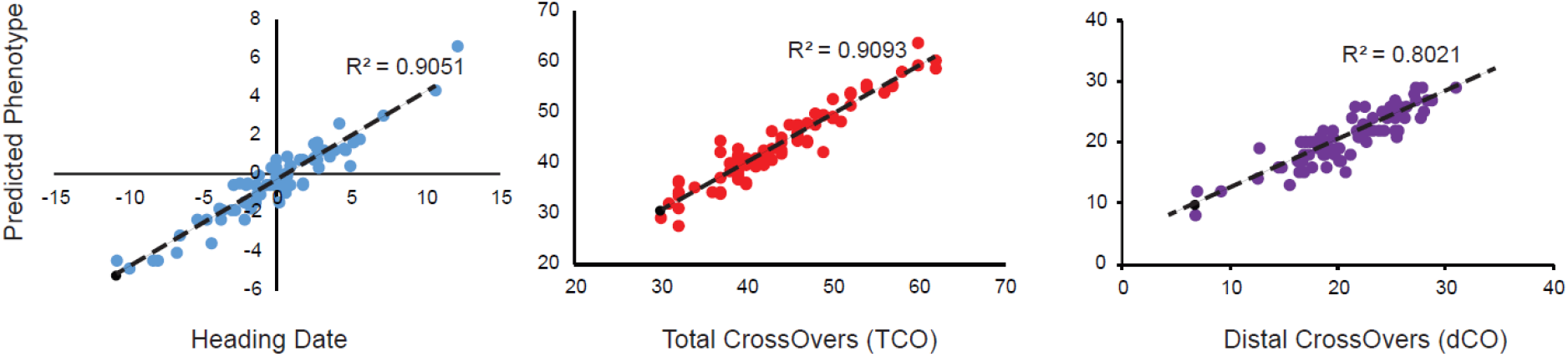
Relationship between the true and predicted phenotypes. Significant markers were identified by stepwise regression on heading date, total numer of crossovers per line (TCO), and total number of distal crossovers per line (dCO) phenotypes.

We also used the imputed genotypes to revisit the genetic analysis of meiotic crossover rate variation in the wheat NAM population (Jordan *et al.* 2018; Blake *et al.* 2019). In the previous study, using a limited number of SNPs genotyped using the 90K iSelect array and GBS, we performed SR analysis and identified 15 and 12 SNPs associated with variation in the total number of crossovers (TCO) and the number of distal crossovers (dCO), respectively (Jordan *et al.* 2018). The identified SNPs explained 48.6% of the variation for TCO and 41% of the variation for dCO. Using the PHG imputed genotypes, we mapped 16 SNPs that together explained 91% of the variance for TCO per line and 12 SNPs explaining 80% of the variance for dCO (Fig. 3, Table S7). Compared to the previous study, SR analyses based on the PHG imputed SNPs detected additional loci with smaller effects on crossover rate (Jordan *et al.* 2018). As a result, the average effect size estimates for TCO and dCO were 2.5 COs and 1.5 COs, respectively. These estimates were lower than the previously reported average effect sizes of 3.36 COs for TCO and 2.3 COs for dCO (Jordan *et al.* 2018). Taken together, these results indicate that the increase in marker density after imputation using the wheat PHG helped to identify new loci with a broader range of effect sizes that together explain a higher proportion of genetic variance compared to the previous study (Jordan *et al.* 2018).

## Discussion

We constructed a wheat PHG database using wheat lines from the major U.S. breeding programs and demonstrated that PHG combined with inexpensive low-coverage genome sequencing could be used to impute genotypes with high accuracy, sufficient to identify variants with smaller effects and support high-resolution mapping studies. Our analyses suggest that the wheat PHG has the potential to effectively utilize community-generated whole-exome capture datasets, currently including thousands of diverse wheat accessions from different geographic regions (Molero *et al.* 2018; He *et al.* 2019; Pont *et al.* 2019), to create a global resource for imputing genotypes. The imputation accuracy provided by the PHG in populations genotyped using the skim-seq, GBS, as well as low-coverage exome sequencing approaches varied, but overall were comparable, indicating that the marker density in the large populations of wheat lines previously genotyped using these methods could be substantially increased by imputation with this newly developed wheat PHG tool.

The accuracy of PHG imputation compared favorably with the commonly used imputation tool Beagle v.5.0 (Browning and Browning 2013), which imputed genotypes with 4% lower accuracy at 0.01x and 0.1x genome coverage than the wheat PHG. In previous studies, imputation of exome capture data with Beagle in populations genotyped using the 90K SNP array and GBS was 93-97% (Jordan *et al.* 2015) and 98% (Nyine *et al.* 2019), respectively. These estimates of accuracy are slightly higher than those obtained in our current study, but overall are comparable, and likely associated with filtering applied to reduce the proportion of missing data in the imputed datasets (Nyine *et al.* 2019), and with the inclusion of more common variants from the array-based genotyping methods. Compared to the imputation accuracy of sorghum (94.1%) and maize (92-95%) PHGs (Jensen *et al.* 2020; Valdes Franco *et al.* 2020), our estimates of accuracy were slightly lower and likely caused by genotyping errors associated with the misalignment of short reads to the more complex, highly repetitive, allopolyploid wheat genome. The higher imputation accuracy in the low-coverage datasets down-sampled from the whole exome capture compared to the accuracy of whole genome skim-seq datasets, which are mostly composed of reads from the repetitive regions of the wheat genome, supports this explanation.

The imputation accuracy among different allele frequency classes improves with an increase in the allele frequency and is higher for a reference allele than for an alternative allele. Consistent with these expectations, the accuracy of imputation in the GBS dataset improved from 87.9% for SNPs with MAF < 0.1 to 92.5% for SNPs with MAF > 0.4, and in the skim-seq dataset from 80.3% for SNPs with MAF < 0.1 to 88.9% for SNPs with MAF >0.4. Previous studies showed that an increase in the reference population size also increases the probability of capturing rare alleles and substantially improves the imputation accuracy of rare variants (Shi *et al.* 2017; Das *et al.* 2018). Our results suggest that the wheat PHG appear to be more effective at utilizing rare haplotypes included into the reference panel for genotype imputation than the commonly used low-coverage imputation method from Beagle. This was demonstarated by imputing genotypes on chromosome 2A, which carries introgression from *Ae. ventricosa* in cultivar Jagger (Cruz *et al.* 2016). The inclusion of genotyping data from cultivar Overley, which also carries this *Ae. ventricosa* introgression, into the PHG database was sufficient for accurate imputation in Jagger. In spite of including genotyping data from cultivar Overley into the reference panel, Beagle imputation of chromosome 2A genotypes in cultivar Jagger was lower compared to PHG. Further efforts aimed at broadening the diversity of accessions in the wheat PHG, including wheat lines carrying known introgressions from wild reatives, will be needed to improve the utility PHG tool for genotype imputation in wheat germplasm.

The application of imputed genotypes to the genetic analyses of trait variation in the wheat NAM population showed that an increase in marker density increases the number of loci associated with trait variation and detects alleles that have smaller effects on phenotypes (*e.g*., recombination rate) than those previously detected using lower density marker sets. The increase in the number of significant loci also resulted in a higher proportion of genetic variance (80-91%) in recombination rate and heading date being explained, suggesting that the imputed genotypes are better at capturing the genetic architecture of these traits, and have the potential to identify more adaptive and beneficial genetic targets in breeding programs.

## Acknowledgements

This project is supported by the Agriculture and Food Research Initiative Competitive Grant 2017–67007-25939 (WheatCAP) from the USDA National Institute of Food and Agriculture, and by the International Wheat Yield Partnership (IWYP). Mention of trade names or commercial products in this publication is solely for the purpose of providing specific information and does not imply recommendation or endorsement by the US Department of Agriculture. USDA is an equal opportunity provider and employee.

## Data availability

The raw sequence data for previously published accessions can be accessed from the NCBI Short-Read Archive database (BioProject SUB2540330 and PRJNA381058). Newly generated exome capture data can be accessed from NCBI Short-Read Archive database (BioProject PRJNA732645). Phenotypic datasets for NAM family 1 associated with the paper can be downloaded from the wheat NAM project website: http://wheatgenomics.plantpath.ksu.edu/nam/.

