## Supplementary Material for "Development of the Wheat Practical Haplotype Graph Database as a Resource for Genotyping Data Storage and Genotype Imputation"

Supplementary Material. Figures S1 and S2; Tables S1-S7.

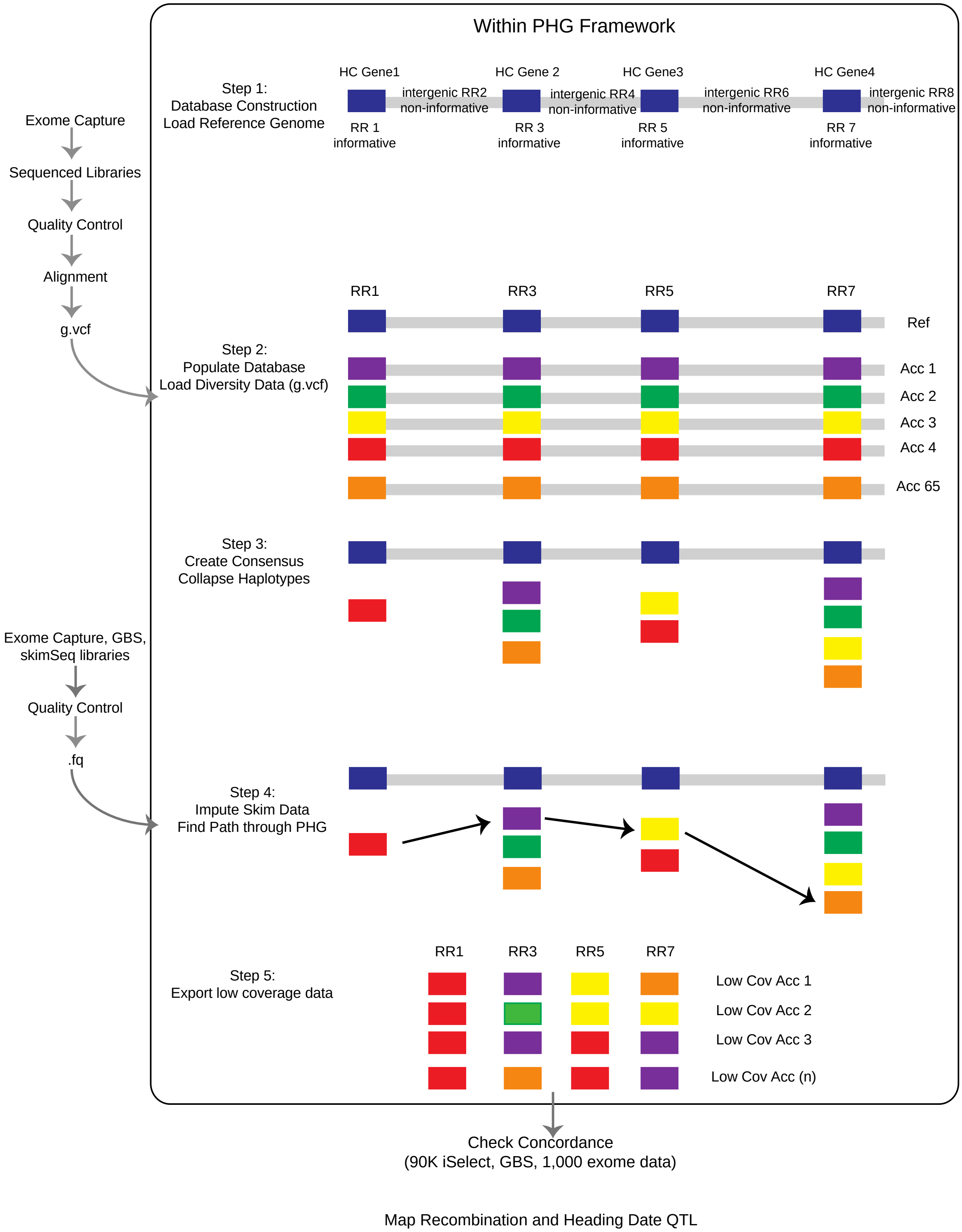


**Figure S1.** The PHG construction workflow.

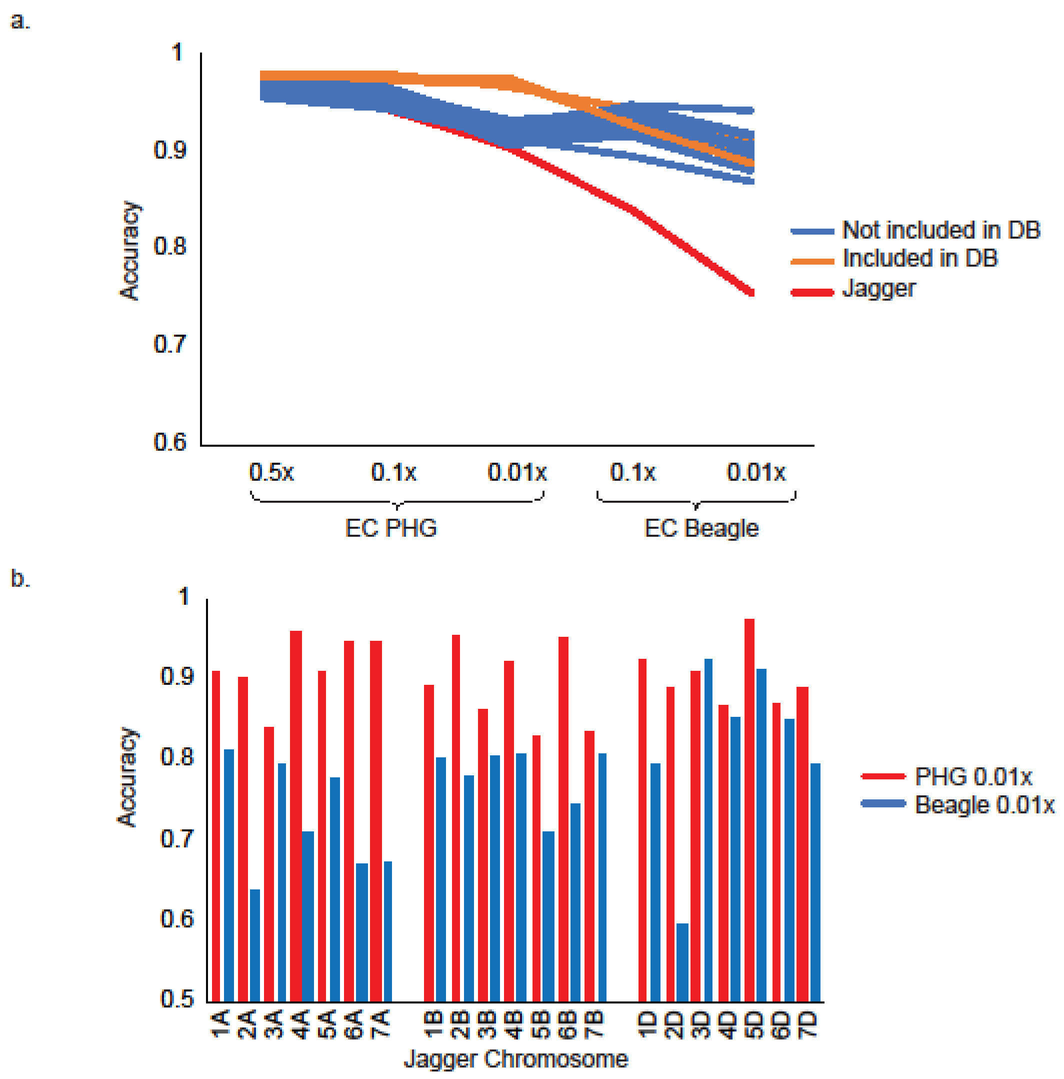


**Figure S2.** Imputation Accuracy Estimates of Wheat Accessions. a) Relationship between imputation accuracy in 20 wheat accessions performed using the Wheat PHG and Beagle and varying depths of sequence read coverage. The sequence data were generated using exome capture (EC). Lines in blue show results for wheat lines not included into the PHG database; orange lines correspond to imputation accuracy obtained for four wheat lines included into the PHG database; the red line represents accuracy of imputation for cultivar Jagger. b) Imputation accuracy estimates for individual wheat chromosomes in cv. Jagger with PHG and Beagle imputation at 0.01x coverage.

**Table S1.** Wheat lines used for the Wheat PHG construction.

| Use | Accession* | WheatCAP Program | NGSC PI# | Origin | Description |
|---|---|---|---|---|---|
| **PHG Database Construction** | 26R61 | Arkansas | PI612153 | Indiana | soft red winter wheat |
| | AGS2000 | Arkansas | PI612956 | Georgia | soft red winter wheat |
| | CO940610* | Colorado State | GSTR 10702 | Colorado | hard winter wheat |
| | Platte* | Colorado State | PI 596297 | Kansas | hard white winter wheat |
| | OPATA | Cornell | PI 591776 | CIMMYT | spring wheat |
| | W7984 (M6) | Cornell | | CIMMYT | synthetic spring wheat |
| | Brundage | Idaho | PI599193, GSTR 13701 | Idaho | soft white winter wheat |
| | Langdon | Idaho | Cltr 13165 | North Dakota | *T. durum* |
| | SY_Capstone | Idaho | PI 665404 | Colorado | hard white spring wheat |
| | UI_Platinum | Idaho | PI 672533 | Idaho | hard white spring Wheat |
| | N87 | Idaho | | | hard spring wheat |
| | MN10201-4-116 | Minnesota | | Minnesota | hard red spring wheat |
| | MN98550-5 | Minnesota | PI660540 | Minnesota | hard red spring wheat |
| | MN99294-1 | Minnesota | | Minnesota | hard red spring wheat |
| | Prosper | Minnesota | PI 662387 | North Dakota | hard red spring wheat |
| | Shelly | Minnesota | PI681618 | Minnesota | hard red spring wheat |
| | Choteau | Montana State University | PI 633974 | Montana | hard red spring wheat |
| | CONAN | Montana State University | PI 607549 | North Dakota | hard red spring wheat |
| | Hank | MSU | PI 613585 | WestBred | hard red spring wheat |
| | LA95135 | NCSU | PI 655291 | Louisiana | soft red winter wheat |
| | SS_mpv57 | NCSU | | Louisiana | soft red winter wheat |
| | BEN | NDSU | | North Dakota | *T. durum* |
| | PI41025 | NDSU | PI41025 | Russia | *T. dicoccum* |
| | Cheyenne* | Nebraska Lincoln | PI 192268 | Nebraska | hard red winter wheat |
| | Billings* | OSU | PI 656843 | Oklahoma | hard red winter wheat |
| | Duster* | OSU, Great Plains | PI 644016 | Oklahoma | hard red winter wheat |
| | TA1615 | SDSU | | WGRC | *Ae. tauschii* |
| | TA1662 | SDSU | PI603230 | | *Ae. tauschii* |
| | TA1718 | SDSU | | WGRC | *Ae. tauschii* |
| | CO960293* | TAMU | | Colorado | hard red winter wheat |
| | TAM111* | TAMU | PI 631352 | Texas | hard red winter wheat |
| | Berkut | CIMMYT | | CIMMYT | spring wheat |
| | CAP2 | CIMMYT | PI 610750 | CIMMYT | synthetic spring wheat |
| | RAC875 | Australia | | Australia | spring wheat |
| | Rio Blanco* | UC Davis | PI531244 | Kansas | hard white winter wheat |
| | Inayama | KSU | PI 382150 | Japan | spring wheat |
| | Bakahtawar94 | none | | Afghanistan | spring wheat |
| | LCS_Star | none | PI673945 | Minnesota Limagrain | hard white spring wheat |
| | Overley* | USDA HWWGRU -KSU | PI 634974 | Kansas | hard red winter wheat |
| | PI634974 | USDA HWWGRU -KSU | PI 634974 | Kansas | hard winter wheat |
| | Overland* | USDA HWWGRU -KSU | PI 647959 | Nebraska | hard winter wheat |
| | Lyman* | USDA HWWGRU -KSU | PI 658067 | South Dakota | hard red winter wheat |
| | Kelse | WSU | PI653842 | Washington | hard red spring wheat |
| | Scarlet | WSU | PI 601814 | Washington | hard red spring wheat |

| | | | | | |
|---|---|---|---|---|---|
| | Altamo | none | | | hard spring wheat |
| | CCW3A37 | none | | | hard winter wheat |
| | CCW3A49 | none | | | hard winter wheat |
| | Dayn | WSU | PI666941 | Washington | hard white spring |
| | IDO444 | none | GSTR 12902 | Idaho | hard winter wheat |
| | KS05HW14-3* | KSU | | Kansas | hard winter wheat |
| | TAM112* | TAMU | PI 643143 | Texas | hard winter wheat |
| | 2045A | NSGC | PI 349512 | Switzerland | spring wheat landrace |
| | Dharwar Dry | none | | India | spring wheat cultivar |
| | PBW343 | none | | India | spring wheat cultivar |
| | Camelot* | Great Plains | PI 653832 | Nebraska | hard red winter wheat |
| | Jagalene* | Great Plains | PI 631376 | Texas | hard red winter wheat |
| | KanMark* | Great Plains | PI 675456 | Kansas | hard red winter wheat |
| | NuPlains* | Great Plains | PI 605741 | Nebraska | hard white winter wheat |
| | Excalibur | none | PI 572701 | Australia | spring wheat |
| | KS061193K-2* | USDA HWWGRU | | Kansas | hard red winter wheat |
| | KS090387K-20* | USDA HWWGRU | | Kansas | hard red winter wheat |
| | WB-Redhawk* | USDA HWWGRU | PI665063 | Kansas | hard red winter wheat |
| | Zenda* | USDA HWWGRU | | Kansas | hard red winter wheat |
| | MOUNTRAIL | USDA, NDSU | PI 607540 | | *T. durum* |
| | MTHW0202 | USDA, NDSU | | | spring wheat |
| **Imputation (Down Sampled)** | Bolles | USDA NDSU | PI678430 | Minnesota | hard red spring wheat |
| | **Duster**** | Great Plains | PI 644016 | Oklahoma | hard red winter wheat |
| | Expedition | Great Plains | PI 629060 | South Dakota | hard red winter wheat |
| | Forefront | USDA HWWGRU | PI 664483 | South Dakota | hard red spring wheat |
| | Goodstreak | USDA HWWGRU | PI 632434 | Nebraska | hard red winter wheat |
| | Ideal | Great Plains | SD05118-1 | South Dakota | hard winter wheat |
| | Jagger | Great Plains | PI 593688 | Kansas | hard red winter wheat |
| | Linkert | USDA NDSU | PI672164 | Minnesota | hard red spring wheat |
| | McGill | USDA HWWGRU | PI 659689 | Nebraska | hard red winter wheat |
| | Mott | USDA NDSU | PI658542 | North Dakota | hard red spring wheat |
| | **NuPlains**** | Great Plains | PI 605741 | Nebraska | hard white winter wheat |
| | **Overley**** | Great Plains | PI 634974 | Kansas | hard red winter wheat |
| | Panhandle | USDA HWWGRU | PI670462 | Nebraska | hard red winter wheat |
| | Prevail | USDA NDSU | PI672486 | South Dakota | hard red spring wheat |
| | Robidoux | USDA HWWGRU | PI 659690 | Nebraska | hard red winter wheat |
| | TAM303 | Great Plains | TX98D1170 | Texas | hard red winter wheat |
| | Traverse | USDA NDSU | PI 642780 | South Dakota | hard red spring wheat |
| | Wesley | Great Plains | PI 605742 | Nebraska | hard red winter wheat |
| | Yellowstone | Great Plains | PI 643428 | Montana | hard red winter wheat |
| | **Zenda**** | USDA HWWGRU | PI 683512 | Kansas | hard red winter wheat |

* lines used in winter wheat IBD analysis
** lines used in PHG database construction

**Table S2.** List of lines subjected to complexity-reduced sequencing.

| Sequencing Technology*/(Population) | ID | Number of Reads* |
|---|---|---|
| **Genotyping by sequencing / (Berkut/Dharwar Dry)** | DHARWAR-1 | 1553300 |
| | DHARWAR-2 | 2127689 |
| | DHARWAR-3 | 2089863 |
| | DHARWAR-4 | 2776216 |
| | DHARWAR-5 | 2110789 |
| | DHARWAR-6 | 1564844 |
| | DHARWAR-7 | 2334733 |
| | DHARWAR-8 | 1698133 |
| | DHARWAR-9 | 1892333 |
| | DHARWAR-10 | 2783586 |
| | DHARWAR-11 | 2440064 |
| | DHARWAR-13 | 2255401 |
| | DHARWAR-15 | 1622381 |
| | DHARWAR-16 | 1375168 |
| | DHARWAR-17 | 1715438 |
| | DHARWAR-18 | 1641959 |
| | DHARWAR-19 | 1871049 |
| | DHARWAR-20 | 2253624 |
| | DHARWAR-21 | 2015967 |
| | DHARWAR-22 | 1951456 |
| | DHARWAR-23 | 1960681 |
| | DHARWAR-24 | 1994041 |
| | DHARWAR-25 | 1590481 |
| | DHARWAR-26 | 1574771 |
| | DHARWAR-27 | 2506973 |
| | DHARWAR-28 | 2433629 |
| | DHARWAR-30 | 2092775 |
| | DHARWAR-32 | 1921740 |
| | DHARWAR-33 | 2495829 |
| | DHARWAR-34 | 1821062 |
| | DHARWAR-35 | 1588191 |
| | DHARWAR-36 | 1288760 |
| | DHARWAR-37 | 1422970 |
| | DHARWAR-38 | 2047759 |
| | DHARWAR-39 | 1750028 |
| | DHARWAR-40 | 2103881 |
| | DHARWAR-41 | 1855342 |
| | DHARWAR-42 | 1797320 |
| | DHARWAR-43 | 2041604 |
| | DHARWAR-44 | 1480090 |
| | DHARWAR-45 | 2566146 |
| | DHARWAR-46 | 2713195 |
| | DHARWAR-48 | 2384553 |
| | DHARWAR-50 | 1948878 |
| | DHARWAR-51 | 1836261 |
| | DHARWAR-52 | 1373420 |
| | DHARWAR-53 | 1672693 |
| | DHARWAR-54 | 1159482 |
| | DHARWAR-55 | 1216116 |

| | | |
|---|---|---|
| | DHARWAR-56 | 1279687 |
| | DHARWAR-57 | 1359086 |
| | DHARWAR-58 | 1482291 |
| | DHARWAR-59 | 1935878 |
| | DHARWAR-60 | 1356332 |
| | DHARWAR-61 | 1678652 |
| | DHARWAR-62 | 1465321 |
| | DHARWAR-63 | 2219051 |
| | DHARWAR-64 | 1307298 |
| | DHARWAR-65 | 1856526 |
| | DHARWAR-66 | 2155352 |
| | DHARWAR-67 | 1785019 |
| | DHARWAR-68 | 1444278 |
| | DHARWAR-70 | 1868376 |
| | DHARWAR-71 | 1325908 |
| | DHARWAR-72 | 1133550 |
| | DHARWAR-73 | 1777633 |
| | DHARWAR-74 | 1899302 |
| | DHARWAR-76 | 1855522 |
| | DHARWAR-77 | 1608282 |
| | DHARWAR-78 | 1815783 |
| | DHARWAR-79 | 2003214 |
| | DHARWAR-80 | 1630167 |
| | DHARWAR-81 | 1943751 |
| | DHARWAR-82 | 1906602 |
| | DHARWAR-83 | 1837817 |
| **Whole Genome Skim-Seq / (Berkut/CI 15144)** | LDRC10-5 | 5809252 |
| | LDRC10-6 | 6041453 |
| | LDRC10-7 | 6442300 |
| | LDRC10-8 | 4554218 |
| | LDRC10-9 | 5928934 |
| | LDRC10-10 | 5051511 |
| | LDRC10-11 | 5037486 |
| | LDRC10-12 | 5158521 |
| | LDRC10-13 | 5219785 |
| | LDRC10-14 | 6774430 |
| | LDRC10-16 | 5580644 |
| | LDRC10-17 | 3212596 |
| | LDRC10-18 | 5094981 |
| | LDRC10-19 | 4159559 |
| | LDRC10-21 | 5072603 |
| | LDRC10-22 | 5669976 |
| | LDRC10-23 | 6366317 |
| | LDRC10-24 | 5991264 |

* GBS reads (1 x 100bp); Whole Genome Skim-Seq (2 x 150bp)

**Table S3. Chromosome Summary of Reference Range and Variants in Wheat PHG**

| Chromosome | Num. reference ranges | All variants | Bi-Allelic SNPs |
|---|---|---|---|
| 1A | 4402 | 64158 | 63302 |
| 1B | 4752 | 83870 | 82890 |
| 1D | 4467 | 67162 | 66513 |
| 2A | 5858 | 77621 | 76692 |
| 2B | 6221 | 103807 | 102428 |
| 2D | 5939 | 108172 | 106906 |
| 3A | 5313 | 54598 | 53956 |
| 3B | 6026 | 77685 | 76833 |
| 3D | 5381 | 80210 | 79406 |
| 4A | 4914 | 59208 | 58389 |
| 4B | 3948 | 32727 | 32482 |
| 4D | 3612 | 39768 | 39575 |
| 5A | 5491 | 48678 | 48268 |
| 5B | 5664 | 70888 | 70191 |
| 5D | 5590 | 78963 | 78345 |
| 6A | 4162 | 56864 | 55988 |
| 6B | 4679 | 77201 | 76165 |
| 6D | 4016 | 63501 | 62813 |
| 7A | 5621 | 74360 | 73455 |
| 7B | 4951 | 63984 | 63271 |
| 7D | 5477 | 90245 | 89453 |
| **Total** | 106,484 | 1,473,670 | 1,457,321 |
| **Average** | 5,071 | 71,175 | 69,396 |

**Table S4. The length of IBD segments (in Mb) shared between the parents of the WheatCAP mapping populations**

| WheatCap Program | Parental Lines | Total IBD Length (Mb) | A genome | B genome | D genome | Mean IBD (Mb)* |
|---|---|---|---|---|---|---|
| Arkansas | 26R61, AGS2000 | 562.72 | 57.70 | 112.57 | 392.45 | 12.79 |
| Cornell | W7986, OPATA | 99.85 | 0 | 0 | 99.85 | 12.48 |
| Colorado | C0940610, Platte | 144.53 | 0 | 19.02 | 125.51 | 9.03 |
| Idaho | SY_Capstone, UI_Platinum | 777.04 | 56.84 | 33.23 | 686.97 | 12.33 |
| Kansas | Overley, Lyman | 128.33 | 15.31 | 16.43 | 96.59 | 10.69 |
| Kansas | Overland, Overley | 273.95 | 9.56 | 12.05 | 252.34 | 11.41 |
| Minnesota | MN98550-5, MN99294-1 | 580.06 | 8.76 | 50.04 | 521.26 | 11.60 |
| Minnesota | Prosper, Shelly | 1287.15 | 265.62 | 286.23 | 735.30 | 11.10 |
| Montana | Hank, Conan | 926.50 | 176.49 | 164.19 | 585.83 | 15.70 |
| North Carolina | SS_mpv57, LA95135 | 632.12 | 16.12 | 115.67 | 500.32 | 11.09 |
| North Dakota | PI41025, Ben | 57.69 | 57.69 | 0 | 0 | 57.69 |
| Oklahoma | Billings, Duster | 435.58 | 43.24 | 2.63 | 389.71 | 11.77 |
| Texas | TAM111, CO960293 | 316.27 | 13.82 | 7.57 | 294.88 | 12.65 |
| California | RAC875, Berkut | 190.24 | 10.86 | 19.85 | 159.54 | 9.51 |
| Washington | Kelse, Scarlet | 359.49 | 75.06 | 55.42 | 229.01 | 14.98 |
| **Average** | | **451.43** | **53.80** | **59.66** | **362.11** | **12.18** |

* average size per region for IBD segments

**Table S5. The length of pairwise IBD segments among winter wheat lines**

| Pairs of lines | Total IBD (Mb) | A genome | B genome | D genome | Mean (Mb) | Proportion of IBD from D genome |
|---|---|---|---|---|---|---|
| Billings CO940610 | 117.77 | 0 | 11.80 | 105.97 | 7.36 | 0.90 |
| Billings Cheyenne | 383.73 | 64.18 | 24.97 | 294.58 | 15.99 | 0.77 |
| Billings CO960293 | 260.68 | 4.37 | 0 | 256.31 | 13.03 | 0.98 |
| Billings KS090387K-20 | 655.74 | 9.70 | 44.50 | 601.54 | 16.39 | 0.92 |
| Billings Lyman | 132.74 | 31.99 | 13.61 | 87.14 | 7.81 | 0.66 |
| Billings NuPlains | 623.56 | 10.80 | 129.37 | 483.39 | 22.27 | 0.78 |
| Billings Platte | 254.10 | 0 | 0 | 254.10 | 14.95 | 1.00 |
| Billings Rio Blanco | 251.81 | 71.22 | 1.99 | 178.59 | 13.25 | 0.71 |
| Billings TAM111 | 260.58 | 65.04 | 33.40 | 162.14 | 11.33 | 0.62 |
| Billings TAM112 | 411.36 | 85.27 | 0 | 326.09 | 8.94 | 0.79 |
| Billings Zenda | 268.57 | 4.23 | 29.64 | 234.70 | 12.21 | 0.87 |
| CO940610 Cheyenne | 286.02 | 162.95 | 15.26 | 107.81 | 16.82 | 0.38 |
| CO940610 CO960293 | 310.29 | 11.83 | 44.74 | 253.72 | 11.08 | 0.82 |
| CO940610 Duster | 117.94 | 15.01 | 0 | 102.93 | 10.72 | 0.87 |
| CO940610 Jagalene | 716.04 | 166.98 | 21.95 | 527.11 | 19.89 | 0.74 |
| CO940610 KS090387K-20 | 381.52 | 183.90 | 15.70 | 181.92 | 16.59 | 0.48 |
| CO940610 Lyman | 299.58 | 12.22 | 22.17 | 265.18 | 9.66 | 0.89 |
| CO940610 NuPlains | 217.90 | 16.74 | 11.26 | 189.90 | 9.90 | 0.87 |
| CO940610 Platte | 144.53 | 0.00 | 19.02 | 125.51 | 9.03 | 0.87 |
| CO940610 Rio Blanco | 121.25 | 11.26 | 19.86 | 90.13 | 6.06 | 0.74 |
| CO940610 TAM111 | 150.91 | 32.34 | 18.14 | 100.43 | 10.06 | 0.67 |
| CO940610 TAM112 | 1237.69 | 58.77 | 106.87 | 1072.05 | 15.87 | 0.87 |
| CO940610 Zenda | 162.27 | 14.21 | 0 | 148.06 | 7.73 | 0.91 |
| Camelot Billings | 299.26 | 4.36 | 7.15 | 287.74 | 12.47 | 0.96 |
| Camelot CO940610 | 280.63 | 13.37 | 0 | 267.26 | 11.23 | 0.95 |
| Camelot Cheyenne | 253.85 | 1.57 | 0 | 252.28 | 9.07 | 0.99 |
| Camelot CO960293 | 450.17 | 156.51 | 3.36 | 290.30 | 16.08 | 0.64 |
| Camelot Duster | 334.15 | 6.34 | 8.53 | 319.29 | 11.14 | 0.96 |
| Camelot Jagalene | 424.34 | 5.60 | 42.29 | 376.45 | 14.63 | 0.89 |
| Camelot KS05HW14-3 | 372.49 | 0 | 42.76 | 329.73 | 13.80 | 0.89 |
| Camelot KS090387K-20 | 551.78 | 20.54 | 154.45 | 376.78 | 20.44 | 0.68 |
| Camelot Lyman | 194.01 | 7.86 | 40.71 | 145.45 | 10.21 | 0.75 |
| Camelot NuPlains | 577.04 | 9.65 | 10.63 | 556.76 | 16.97 | 0.96 |
| Camelot Platte | 289.94 | 0 | 31.51 | 258.43 | 13.18 | 0.89 |
| Camelot Rio Blanco | 312.95 | 0 | 20.46 | 292.49 | 13.04 | 0.93 |
| Camelot TAM111 | 208.99 | 11.85 | 9.84 | 187.30 | 11.00 | 0.90 |
| Camelot TAM112 | 246.39 | 4.16 | 9.51 | 232.72 | 7.70 | 0.94 |
| Camelot Zenda | 212.16 | 12.86 | 21.26 | 178.04 | 8.49 | 0.84 |
| Cheyenne CO960293 | 367.67 | 0 | 4.72 | 362.95 | 13.62 | 0.99 |

| | | | | | | |
|---|---|---|---|---|---|---|
| Cheyenne Duster | 292.85 | 3.77 | 21.09 | 267.99 | 13.31 | 0.92 |
| Cheyenne Lyman | 368.10 | 8.72 | 62.91 | 296.47 | 11.50 | 0.81 |
| Cheyenne Rio Blanco | 276.29 | 3.22 | 35.31 | 237.75 | 8.91 | 0.86 |
| Cheyenne Zenda | 321.27 | 0 | 19.85 | 301.42 | 16.06 | 0.94 |
| Duster CO960293 | 160.50 | 7.19 | 0 | 153.31 | 7.30 | 0.96 |
| Duster Lyman | 301.04 | 2.72 | 45.77 | 252.55 | 15.84 | 0.84 |
| Duster Rio Blanco | 226.59 | 21.27 | 8.12 | 197.19 | 9.85 | 0.87 |
| Duster Zenda | 239.10 | 4.91 | 24.59 | 209.60 | 11.96 | 0.88 |
| Jagalene Cheyenne | 463.78 | 3.24 | 123.57 | 336.97 | 15.46 | 0.73 |
| Jagalene CO960293 | 350.50 | 39.11 | 4.10 | 307.30 | 15.24 | 0.88 |
| Jagalene Duster | 325.72 | 0 | 153.14 | 172.58 | 11.23 | 0.53 |
| Jagalene Lyman | 211.65 | 0 | 58.56 | 153.09 | 11.14 | 0.72 |
| Jagalene Rio Blanco | 376.68 | 17.92 | 70.91 | 287.85 | 11.77 | 0.76 |
| Jagalene Zenda | 530.44 | 135.81 | 51.02 | 343.62 | 17.11 | 0.65 |
| KanMark Billings | 245.78 | 24.56 | 8.01 | 213.21 | 8.78 | 0.87 |
| KanMark CO940610 | 255.83 | 2.32 | 6.99 | 246.52 | 13.46 | 0.96 |
| KanMark Camelot | 269.75 | 10.05 | 3.54 | 256.16 | 12.26 | 0.95 |
| KanMark Cheyenne | 325.95 | 7.29 | 86.78 | 231.88 | 12.07 | 0.71 |
| KanMark CO960293 | 412.39 | 23.12 | 237.62 | 151.65 | 17.93 | 0.37 |
| KanMark Duster | 563.43 | 8.78 | 37.81 | 516.84 | 14.09 | 0.92 |
| KanMark Jagalene | 660.75 | 185.06 | 219.39 | 256.30 | 19.43 | 0.39 |
| KanMark KS05HW14-3 | 526.02 | 161.90 | 144.61 | 219.51 | 18.14 | 0.42 |
| KanMark KS090387K-20 | 648.87 | 27.80 | 137.88 | 483.18 | 16.22 | 0.74 |
| KanMark Lyman | 295.13 | 2.50 | 39.29 | 253.34 | 13.41 | 0.86 |
| KanMark NuPlains | 533.09 | 68.89 | 199.66 | 264.54 | 13.67 | 0.50 |
| KanMark Platte | 374.03 | 63.49 | 44.97 | 265.56 | 12.90 | 0.71 |
| KanMark Rio Blanco | 357.22 | 28.45 | 26.20 | 302.57 | 11.52 | 0.85 |
| KanMark TAM111 | 473.65 | 113.06 | 69.95 | 290.63 | 18.22 | 0.61 |
| KanMark TAM112 | 783.11 | 110.57 | 132.23 | 540.31 | 12.24 | 0.69 |
| KanMark Zenda | 358.53 | 13.37 | 27.82 | 317.34 | 11.95 | 0.89 |
| KS05HW14-3 Billings | 363.19 | 1.80 | 32.02 | 329.36 | 12.11 | 0.91 |
| KS05HW14-3 CO940610 | 385.50 | 8.87 | 52.62 | 324.02 | 10.14 | 0.84 |
| KS05HW14-3 Cheyenne | 409.63 | 5.89 | 4.89 | 398.85 | 11.07 | 0.97 |
| KS05HW14-3 CO960293 | 638.99 | 64.92 | 26.74 | 547.33 | 11.41 | 0.86 |
| KS05HW14-3 Duster | 292.18 | 0 | 2.07 | 290.11 | 11.69 | 0.99 |
| KS05HW14-3 Jagalene | 373.48 | 25.24 | 0 | 348.24 | 14.94 | 0.93 |
| KS05HW14-3 KS090387K-20 | 204.89 | 0 | 39.29 | 165.60 | 8.20 | 0.81 |
| KS05HW14-3 Lyman | 197.02 | 6.51 | 10.10 | 180.41 | 8.57 | 0.92 |
| KS05HW14-3 NuPlains | 602.62 | 17.33 | 16.05 | 569.24 | 17.22 | 0.94 |
| KS05HW14-3 Platte | 396.77 | 0 | 10.36 | 386.41 | 18.04 | 0.97 |
| KS05HW14-3 Rio Blanco | 552.22 | 3.07 | 31.36 | 517.79 | 14.16 | 0.94 |
| KS05HW14-3 TAM111 | 499.99 | 0 | 48.10 | 451.89 | 20.00 | 0.90 |
| KS05HW14-3 TAM112 | 944.70 | 162.53 | 21.95 | 760.22 | 13.89 | 0.80 |

| | | | | | | |
|---|---|---|---|---|---|---|
| KS05HW14-3 Zenda | 538.49 | 29.81 | 100.78 | 407.90 | 16.32 | 0.76 |
| KS061193K-2 Billings | 333.73 | 27.81 | 3.32 | 302.60 | 13.35 | 0.91 |
| KS061193K-2 CO940610 | 542.40 | 0 | 12.73 | 529.66 | 16.95 | 0.98 |
| KS061193K-2 Camelot | 672.79 | 11.83 | 232.40 | 428.57 | 18.18 | 0.64 |
| KS061193K-2 Cheyenne | 214.80 | 1.90 | 74.15 | 138.75 | 11.93 | 0.65 |
| KS061193K-2 CO960293 | 270.22 | 26.13 | 4.78 | 239.31 | 12.28 | 0.89 |
| KS061193K-2 Duster | 408.25 | 52.81 | 28.80 | 326.64 | 15.12 | 0.80 |
| KS061193K-2 Jagalene | 925.45 | 37.28 | 70.51 | 817.66 | 17.14 | 0.88 |
| KS061193K-2 KanMark | 536.37 | 34.97 | 18.67 | 482.72 | 13.41 | 0.90 |
| KS061193K-2 KS05HW14-3 | 199.26 | 8.55 | 6.95 | 183.75 | 8.30 | 0.92 |
| KS061193K-2 KS090387K-20 | 888.91 | 4.92 | 123.41 | 760.57 | 18.14 | 0.86 |
| KS061193K-2 Lyman | 233.11 | 0 | 37.95 | 195.17 | 14.57 | 0.84 |
| KS061193K-2 NuPlains | 365.28 | 20.30 | 10.55 | 334.44 | 14.05 | 0.92 |
| KS061193K-2 Platte | 86.58 | 0 | 13.52 | 73.06 | 5.41 | 0.84 |
| KS061193K-2 Rio Blanco | 235.99 | 13.07 | 7.83 | 215.08 | 11.24 | 0.91 |
| KS061193K-2 TAM111 | 231.02 | 0 | 21.24 | 209.78 | 14.44 | 0.91 |
| KS061193K-2 TAM112 | 601.47 | 128.80 | 4.06 | 468.61 | 13.08 | 0.78 |
| KS061193K-2 Zenda | 900.67 | 153.16 | 74.06 | 673.45 | 13.86 | 0.75 |
| KS090387K-20 Cheyenne | 232.68 | 14.80 | 4.53 | 213.36 | 11.63 | 0.92 |
| KS090387K-20 CO960293 | 162.74 | 9.18 | 22.52 | 131.04 | 8.57 | 0.81 |
| KS090387K-20 Duster | 524.78 | 1.51 | 16.05 | 507.22 | 20.18 | 0.97 |
| KS090387K-20 Jagalene | 773.29 | 160.99 | 24.50 | 587.80 | 19.83 | 0.76 |
| KS090387K-20 Lyman | 141.97 | 0.00 | 23.78 | 118.19 | 10.14 | 0.83 |
| KS090387K-20 Rio Blanco | 146.21 | 36.35 | 2.99 | 106.86 | 9.14 | 0.73 |
| KS090387K-20 Zenda | 563.39 | 7.81 | 6.01 | 549.56 | 18.17 | 0.98 |
| Lyman CO960293 | 301.78 | 0 | 11.57 | 290.20 | 12.07 | 0.96 |
| NuPlains Cheyenne | 352.63 | 0 | 18.20 | 334.43 | 13.06 | 0.95 |
| NuPlains CO960293 | 360.12 | 0 | 21.21 | 338.90 | 18.95 | 0.94 |
| NuPlains Duster | 346.44 | 0 | 0 | 346.44 | 14.44 | 1.00 |
| NuPlains Jagalene | 592.13 | 0 | 187.06 | 405.07 | 14.44 | 0.68 |
| NuPlains KS090387K-20 | 484.73 | 0 | 43.58 | 441.15 | 16.71 | 0.91 |
| NuPlains Lyman | 381.45 | 182.79 | 12.75 | 185.91 | 18.16 | 0.49 |
| NuPlains Platte | 612.13 | 13.58 | 0 | 598.55 | 16.54 | 0.98 |
| NuPlains Rio Blanco | 596.49 | 6.42 | 12.92 | 577.15 | 11.93 | 0.97 |
| NuPlains Zenda | 605.73 | 166.82 | 40.12 | 398.79 | 14.42 | 0.66 |
| Overland Billings | 236.61 | 5.39 | 0 | 231.22 | 11.83 | 0.98 |
| Overland CO940610 | 250.61 | 0 | 25.54 | 225.07 | 9.64 | 0.90 |
| Overland Camelot | 271.77 | 8.75 | 28.86 | 234.16 | 9.71 | 0.86 |
| Overland Cheyenne | 411.11 | 8.37 | 17.43 | 385.30 | 9.79 | 0.94 |
| Overland CO960293 | 325.30 | 57.27 | 4.61 | 263.42 | 10.49 | 0.81 |
| Overland Duster | 260.06 | 1.92 | 19.49 | 238.66 | 12.38 | 0.92 |
| Overland Jagalene | 231.81 | 0 | 19.64 | 212.17 | 13.64 | 0.92 |
| Overland KanMark | 327.19 | 129.11 | 6.14 | 191.94 | 12.12 | 0.59 |

| | | | | | | |
|---|---|---|---|---|---|---|
| Overland KS05HW14-3 | 496.82 | 13.75 | 16.34 | 466.73 | 11.83 | 0.94 |
| Overland KS061193K-2 | 279.06 | 12.20 | 20.28 | 246.58 | 9.00 | 0.88 |
| Overland KS090387K-20 | 337.49 | 19.35 | 82.00 | 236.14 | 12.05 | 0.70 |
| Overland Lyman | 305.67 | 20.53 | 34.90 | 250.24 | 8.49 | 0.82 |
| Overland NuPlains | 409.18 | 3.53 | 4.15 | 401.50 | 15.74 | 0.98 |
| Overland Platte | 154.59 | 0 | 12.11 | 142.49 | 11.04 | 0.92 |
| Overland Rio Blanco | 285.18 | 17.31 | 34.83 | 233.04 | 8.91 | 0.82 |
| Overland TAM111 | 298.53 | 0 | 24.40 | 274.13 | 12.98 | 0.92 |
| Overland TAM112 | 530.96 | 107.39 | 0 | 423.57 | 13.97 | 0.80 |
| Overland WB-Redhawk | 335.09 | 0 | 12.69 | 322.40 | 16.75 | 0.96 |
| Overland Zenda | 219.93 | 5.66 | 12.11 | 202.15 | 8.46 | 0.92 |
| Overley Billings | 359.95 | 13.31 | 3.32 | 343.31 | 8.57 | 0.95 |
| Overley Camelot | 586.57 | 0 | 20.29 | 566.29 | 12.75 | 0.97 |
| Overley Cheyenne | 145.52 | 0 | 23.15 | 122.37 | 7.28 | 0.84 |
| Overley CO960293 | 1337.21 | 466.99 | 309.00 | 561.23 | 22.29 | 0.42 |
| Overley Duster | 289.37 | 0 | 34.03 | 255.34 | 6.58 | 0.88 |
| Overley Jagalene | 1724.39 | 155.80 | 49.21 | 1519.38 | 14.37 | 0.88 |
| Overley KanMark | 346.16 | 126.05 | 7.13 | 212.98 | 9.62 | 0.62 |
| Overley KS05HW14-3 | 810.58 | 5.18 | 103.16 | 702.24 | 13.07 | 0.87 |
| Overley KS061193K-2 | 1371.07 | 77.34 | 83.27 | 1210.46 | 13.44 | 0.88 |
| Overley KS090387K-20 | 710.75 | 112.71 | 17.52 | 580.52 | 12.69 | 0.82 |
| Overley Lyman | 253.56 | 0 | 0 | 253.56 | 14.09 | 1.00 |
| Overley NuPlains | 310.20 | 8.64 | 82.79 | 218.77 | 8.62 | 0.71 |
| Overley Overland | 274.77 | 3.20 | 26.20 | 245.37 | 7.23 | 0.89 |
| Overley Platte | 386.24 | 0 | 199.35 | 186.90 | 12.07 | 0.48 |
| Overley Rio Blanco | 494.16 | 27.42 | 3.88 | 462.87 | 16.47 | 0.94 |
| Overley TAM111 | 200.59 | 8.84 | 46.97 | 144.77 | 7.71 | 0.72 |
| Overley TAM112 | 1474.22 | 69.45 | 65.95 | 1338.82 | 12.93 | 0.91 |
| Overley WB-Redhawk | 1282.80 | 84.99 | 226.85 | 970.96 | 11.45 | 0.76 |
| Overley Zenda | 962.63 | 29.00 | 282.69 | 650.94 | 10.46 | 0.68 |
| Platte Cheyenne | 350.54 | 5.08 | 13.89 | 331.57 | 19.47 | 0.95 |
| Platte CO960293 | 543.94 | 0 | 116.24 | 427.70 | 19.43 | 0.79 |
| Platte Duster | 362.64 | 0 | 140.26 | 222.38 | 12.95 | 0.61 |
| Platte Jagalene | 523.43 | 2.25 | 232.38 | 288.80 | 10.68 | 0.55 |
| Platte KS090387K-20 | 200.51 | 88.67 | 2.66 | 109.17 | 11.14 | 0.54 |
| Platte Lyman | 94.75 | 0 | 14.14 | 80.61 | 13.54 | 0.85 |
| Platte Rio Blanco | 524.30 | 3.46 | 15.06 | 505.77 | 12.79 | 0.96 |
| Platte Zenda | 164.69 | 0 | 17.86 | 146.83 | 11.76 | 0.89 |
| RioBlanco CO960293 | 353.35 | 19.40 | 8.36 | 325.59 | 12.18 | 0.92 |
| RioBlanco Lyman | 328.26 | 2.95 | 35.88 | 289.43 | 13.68 | 0.88 |
| RioBlanco Zenda | 146.01 | 17.46 | 4.64 | 123.91 | 6.35 | 0.85 |
| TAM111 Cheyenne | 341.39 | 6.05 | 26.86 | 308.48 | 12.64 | 0.90 |
| TAM111 CO960293 | 316.27 | 13.82 | 7.57 | 294.88 | 12.65 | 0.93 |

| | | | | | | |
|---|---|---|---|---|---|---|
| TAM111 Duster | 389.74 | 14.78 | 13.98 | 360.98 | 14.43 | 0.93 |
| TAM111 Jagalene | 277.96 | 0 | 9.91 | 268.05 | 21.38 | 0.96 |
| TAM111 KS090387K-20 | 332.52 | 6.52 | 4.31 | 321.69 | 16.63 | 0.97 |
| TAM111 Lyman | 175.23 | 60.05 | 0 | 115.18 | 14.60 | 0.66 |
| TAM111 NuPlains | 523.40 | 82.01 | 89.27 | 352.13 | 18.69 | 0.67 |
| TAM111 Platte | 141.50 | 18.67 | 9.91 | 112.91 | 10.11 | 0.80 |
| TAM111 Rio Blanco | 469.35 | 8.36 | 6.71 | 454.28 | 15.64 | 0.97 |
| TAM111 Zenda | 188.03 | 11.55 | 22.01 | 154.47 | 12.54 | 0.82 |
| TAM112 Cheyenne | 525.87 | 8.21 | 5.47 | 512.19 | 14.61 | 0.97 |
| TAM112 CO960293 | 630.70 | 103.61 | 60.53 | 466.56 | 12.61 | 0.74 |
| TAM112 Duster | 501.75 | 57.37 | 10.03 | 434.36 | 10.04 | 0.87 |
| TAM112 Jagalene | 809.44 | 5.79 | 59.50 | 744.16 | 16.86 | 0.92 |
| TAM112 KS090387K-20 | 823.87 | 20.79 | 61.40 | 741.68 | 15.26 | 0.90 |
| TAM112 Lyman | 423.22 | 30.21 | 24.32 | 368.69 | 8.14 | 0.87 |
| TAM112 NuPlains | 790.38 | 287.69 | 37.76 | 464.93 | 10.98 | 0.59 |
| TAM112 Platte | 336.91 | 3.26 | 14.16 | 319.49 | 12.96 | 0.95 |
| TAM112 Rio Blanco | 510.22 | 110.29 | 28.10 | 371.83 | 13.43 | 0.73 |
| TAM112 TAM111 | 856.76 | 15.44 | 83.65 | 757.68 | 12.24 | 0.88 |
| TAM112 Zenda | 403.83 | 64.15 | 7.95 | 331.73 | 8.78 | 0.82 |
| WB-Redhawk Billings | 257.26 | 21.72 | 3.92 | 231.62 | 13.54 | 0.90 |
| WB-Redhawk CO940610 | 354.51 | 24.12 | 51.95 | 278.44 | 12.66 | 0.79 |
| WB-Redhawk Camelot | 664.79 | 114.01 | 7.55 | 543.24 | 22.92 | 0.82 |
| WB-Redhawk Cheyenne | 119.20 | 15.02 | 40.81 | 63.36 | 7.95 | 0.53 |
| WB-Redhawk CO960293 | 202.23 | 30.84 | 64.20 | 107.19 | 11.23 | 0.53 |
| WB-Redhawk Duster | 311.92 | 0 | 10.28 | 301.64 | 13.00 | 0.97 |
| WB-Redhawk Jagalene | 754.22 | 40.32 | 149.41 | 564.49 | 18.40 | 0.75 |
| WB-Redhawk KanMark | 310.19 | 25.53 | 36.43 | 248.24 | 10.34 | 0.80 |
| WB-Redhawk KS05HW14-3 | 459.66 | 8.08 | 6.54 | 445.05 | 14.36 | 0.97 |
| WB-Redhawk KS061193K-2 | 784.86 | 135.43 | 34.89 | 614.55 | 15.70 | 0.78 |
| WB-Redhawk KS090387K-20 | 437.75 | 9.40 | 41.06 | 387.29 | 11.22 | 0.88 |
| WB-Redhawk Lyman | 164.48 | 21.74 | 10.79 | 131.96 | 10.97 | 0.80 |
| WB-Redhawk NuPlains | 606.29 | 27.18 | 16.68 | 562.43 | 16.84 | 0.93 |
| WB-Redhawk Platte | 211.39 | 7.28 | 15.46 | 188.65 | 11.74 | 0.89 |
| WB-Redhawk Rio Blanco | 375.12 | 27.53 | 156.98 | 190.61 | 17.86 | 0.51 |
| WB-Redhawk TAM111 | 184.21 | 0 | 44.21 | 139.99 | 9.70 | 0.76 |
| WB-Redhawk TAM112 | 615.78 | 0 | 49.47 | 566.30 | 9.93 | 0.92 |
| WB-Redhawk Zenda | 615.34 | 15.12 | 57.23 | 542.99 | 12.82 | 0.88 |
| Zenda CO960293 | 244.21 | 15.07 | 0 | 229.14 | 11.63 | 0.94 |
| Zenda Lyman | 191.35 | 0 | 0 | 191.35 | 13.67 | 1.00 |
| **Average** | **414.96** | **33.73** | **41.27** | **339.96** | **12.99** | **0.83** |

Table S6. Imputation Accuracy for different methods of target population genotyping (GBS and Skim-seq).

| Method of genotyping | Line ID | Accuracy |
|---|---|---|
| GBS | DHARWAR-1 | 0.9072 |
| | DHARWAR-2 | 0.9023 |
| | DHARWAR-3 | 0.9022 |
| | DHARWAR-4 | 0.9004 |
| | DHARWAR-5 | 0.8921 |
| | DHARWAR-6 | 0.9112 |
| | DHARWAR-7 | 0.9009 |
| | DHARWAR-8 | 0.8993 |
| | DHARWAR-9 | 0.9016 |
| | DHARWAR-10 | 0.8966 |
| | DHARWAR-11 | 0.9051 |
| | DHARWAR-13 | 0.9038 |
| | DHARWAR-15 | 0.9061 |
| | DHARWAR-16 | 0.9065 |
| | DHARWAR-17 | 0.9049 |
| | DHARWAR-18 | 0.9053 |
| | DHARWAR-19 | 0.9005 |
| | DHARWAR-20 | 0.9053 |
| | DHARWAR-21 | 0.9103 |
| | DHARWAR-22 | 0.9106 |
| | DHARWAR-23 | 0.9028 |
| | DHARWAR-24 | 0.9019 |
| | DHARWAR-25 | 0.9033 |
| | DHARWAR-26 | 0.9059 |
| | DHARWAR-27 | 0.9007 |
| | DHARWAR-28 | 0.9042 |
| | DHARWAR-30 | 0.903 |
| | DHARWAR-32 | 0.909 |
| | DHARWAR-33 | 0.8953 |
| | DHARWAR-34 | 0.9054 |
| | DHARWAR-35 | 0.9038 |
| | DHARWAR-36 | 0.9025 |
| | DHARWAR-37 | 0.9096 |
| | DHARWAR-38 | 0.9005 |
| | DHARWAR-39 | 0.9144 |
| | DHARWAR-40 | 0.9031 |
| | DHARWAR-41 | 0.9076 |
| | DHARWAR-42 | 0.9101 |
| | DHARWAR-43 | 0.9055 |

| | | |
|---|---|---|
| | DHARWAR-44 | 0.9062 |
| | DHARWAR-45 | 0.9 |
| | DHARWAR-46 | 0.9027 |
| | DHARWAR-48 | 0.8972 |
| | DHARWAR-50 | 0.9056 |
| | DHARWAR-51 | 0.9022 |
| | DHARWAR-52 | 0.9131 |
| | DHARWAR-53 | 0.9012 |
| | DHARWAR-54 | 0.907 |
| | DHARWAR-55 | 0.9131 |
| | DHARWAR-56 | 0.9073 |
| | DHARWAR-57 | 0.9057 |
| | DHARWAR-58 | 0.8998 |
| | DHARWAR-59 | 0.9057 |
| | DHARWAR-60 | 0.8991 |
| | DHARWAR-61 | 0.9093 |
| | DHARWAR-62 | 0.9089 |
| | DHARWAR-63 | 0.9045 |
| | DHARWAR-64 | 0.9007 |
| | DHARWAR-65 | 0.9036 |
| | DHARWAR-66 | 0.9003 |
| | DHARWAR-67 | 0.9028 |
| | DHARWAR-68 | 0.9077 |
| | DHARWAR-70 | 0.9075 |
| | DHARWAR-71 | 0.9088 |
| | DHARWAR-72 | 0.9049 |
| | DHARWAR-73 | 0.9047 |
| | DHARWAR-74 | 0.9045 |
| | DHARWAR-76 | 0.9027 |
| | DHARWAR-77 | 0.9046 |
| | DHARWAR-78 | 0.9077 |
| | DHARWAR-79 | 0.9 |
| | DHARWAR-80 | 0.9068 |
| | DHARWAR-81 | 0.9058 |
| | DHARWAR-82 | 0.9031 |
| | DHARWAR-83 | 0.8903 |
| | **Average** | **0.9042** |
| **skim Seq** | LDRC10-5 | 0.8671 |
| | LDRC10-6 | 0.8573 |
| | LDRC10-7 | 0.8499 |
| | LDRC10-8 | 0.8514 |
| | LDRC10-9 | 0.8644 |
| | LDRC10-10 | 0.8467 |

| | | |
|---|---|---|
| | LDRC10-11 | 0.8583 |
| | LDRC10-12 | 0.8487 |
| | LDRC10-13 | 0.8646 |
| | LDRC10-14 | 0.8661 |
| | LDRC10-16 | 0.8466 |
| | LDRC10-17 | 0.8217 |
| | LDRC10-18 | 0.8419 |
| | LDRC10-19 | 0.8428 |
| | LDRC10-21 | 0.8415 |
| | LDRC10-22 | 0.8532 |
| | LDRC10-23 | 0.8750 |
| | LDRC10-24 | 0.8539 |
| | **Average** | **0.8528** |

**Table S7. Regression Analysis for NAM1 phenotypes**

| Trait | Chromosome | Position, bp | Effect* | $R^2$ |
|---|---|---|---|---|
| **TCO**** | 1A | 24338410 | 2.8561 | 90.9 |
| | 2A | 690924031 | -1.0464 | |
| | 2B | 655213027 | 1.224 | |
| | 2B | 681349687 | -5.4315 | |
| | 2D | 641109363 | -5.3329 | |
| | 3A | 8316383 | 4.2101 | |
| | 3A | 13536555 | -2.0524 | |
| | 3A | 24038281 | -2.2197 | |
| | 3A | 574459200 | -2.1884 | |
| | 3A | 732911987 | -2.1622 | |
| | 5B | 564988417 | -2.1359 | |
| | 5D | 41604006 | 1.5155 | |
| | 5D | 562723413 | 1.6485 | |
| | 6B | 15932507 | 1.1151 | |
| | 6D | 346977546 | 2.61 | |
| | 7A | 708319323 | 2.17 | |
| **HD**** | 1B | 62419179 | -0.5365 | 90.5 |
| | 1D | 42700817 | 0.4259 | |
| | 2A | 142314578 | -0.7482 | |
| | 2B | 742815469 | 0.9514 | |
| | 2B | 800188720 | 0.6988 | |
| | 3A | 649632776 | -0.4625 | |
| | 3B | 580786270 | 0.5427 | |
| | 3D | 525536813 | -0.6617 | |
| | 4A | 95378425 | -0.7689 | |
| | 5A | 580794434 | -0.79765 | |
| | 5D | 542110241 | -0.57484 | |

*Allelic estimates represent effect for Chinese Spring reference allele
** HD – heading date; TCO – total number of crossovers.
